# Cell-to-cell heterogeneity in Sox2 and Brachyury expression ratios guides progenitor destiny by controlling their motility.

**DOI:** 10.1101/2020.11.18.388611

**Authors:** Michèle Romanos, Guillaume Allio, Léa Combres, Francois Médevielle, Nathalie Escalas, Cathy Soula, Ben Steventon, Ariane Trescases, Bertrand Bénazéraf

## Abstract

Although cell-to-cell heterogeneity in gene and protein expression within cell populations has been widely documented, we know little about its potential biological functions. We addressed this issue by studying progenitors that populate the posterior region of the vertebrate embryos, a cell population known for its capacity to self-renew or to contribute to the formation of the neural tube and paraxial mesoderm tissues. Posterior progenitors are characterized by the co-expression of Sox2 and Brachyury (Bra), two transcription factors related to neural and mesodermal lineages, respectively. In this study, we show that the respective levels of Sox2 and Bra proteins display a high degree of variability among posterior progenitors of the quail embryo. By developing forced expression and downregulation approaches, we further provide evidence that the value of the Sox2-to-Bra ratio in a given progenitor directs its choice of staying in place or exit the progenitor zone to generate neural or mesodermal cells. Time-lapse imaging together with mathematical modeling then reveal that variations of the Sox2-to-Bra ratio confer these cells heterogeneous motile behaviors. While high Bra levels display high motile properties that push cells to join the mesoderm, high levels of Sox2 tend to inhibit cell movement making cells get integrated into the neural tube. Our work thus provides evidence that the spatial heterogeneity of posterior progenitors, with regards to their relative contents of Sox2 and Bra and thus to their motile properties, is fundamental to maintain a pool of resident progenitors while others segregate to contribute to tissue formation. More broadly, our work reveals that heterogeneity among a population of progenitor cells is critical to ensure robust multi-tissue morphogenesis.

## Introduction

Cells are the functional units of living organisms. During embryogenesis, they divide and specify in multiple cell types that organize spatially into tissues and organs. Specification events take place under the influences of the cell’s own history and of environmental clues. Over the last years, access to new technologies has revealed that embryonic cells often display an unappreciated level of heterogeneity. For instance, gene expression analyses suggest that, within the same embryonic tissue, cells which were thought to be either equivalent or different, are actually organized into a continuum of various specification states (1,2). The impact of this new level of complexity on morphogenesis has not been extensively explored due to the difficulty of experimentally manipulating expression levels within targeted populations of cells *in vivo*. Progenitor cells located at the posterior tip of the vertebrate embryo, in an area known as the progenitor zone (PZ), constitute a great model to study how a population of stem-like cells develops into different cell types. The use of fluorescent tracers in bird and mouse embryos has revealed that cells of the PZ, called here posterior progenitors, contribute to formation of the presomitic mesoderm (PSM), the mesodermal tissue that generates muscle and vertebrae but also of the neural tube (NT), the neuro-ectodermal tissue that gives rise to the central nervous system (3–6). These studies also evidenced different cell behaviors with some cells exiting the PZ and others remaining resident in this area. Grafting experiments next showed that resident posterior progenitors have the capacity to self-renew while providing new neural and mesodermal progenies (7,8), thus indicating that the PZ contains different types of progenitors. Heterogeneity of the posterior progenitor population was further confirmed by retrospective clonal analysis studies performed in the mouse embryo which revealed the existence of single progenitors, giving rise either to neural or mesodermal cells, but also of bi-potent progenitors, named neuro-mesodermal progenitors (NMPs), that generate both neural and mesodermal cells (9). The existence of bi-potent progenitors has since been shown at earlier stages of zebrafish development (10) and in bird embryos (11–13). Thus, to sustain formation of tissues that compose the vertebrate body axis, the heterogeneous population of posterior progenitors must maintain an appropriate balance between the two choices of staying in place and self-renew or exit the progenitor region to contribute to formation of the NT and the PSM. How this balance is established and controlled over time remains an open question.

Two transcription factors, Sox2 (SRY sex determining region Y-box 2) and Bra (Brachyury), have been described for their respective roles in neural and mesodermal specification during embryonic development (14,15). Sox2 is known to be expressed in the neural progenitors that form the neural tube where it contributes to maintain their undifferentiated state. Its involvement in neural specification has also been revealed by a study showing that ectopic expression of Sox2 in cells of the PSM is sufficient to reprogram these cells, which then adopt a neural identity (16). Bra protein was initially identified for its essential function in formation of the paraxial mesoderm during the posterior extension phase (14,17). Its crucial role in mesodermal specification has been demonstrated, in particular, by phenotypic study of chimeric mouse embryos composed of both Bra mutant and wild-type cells, and in which only wild-type cells are capable of generating mesodermal cells (18). More recent studies have shown that Sox2 and Bra are expressed in posterior progenitors of developing embryos, indicating that activation of their expression takes place in progenitor cells before these cells colonize the NT or the PSM (19–21). In addition, these studies have shown that both proteins are co-expressed in progenitor cells, an observation consistent with the presence of bi-potent progenitors in this tissue. Works done in mouse embryo and in *in vitro* systems derived from embryonic stem cells indicate that Bra and Sox2 influence the choice between neural and mesodermal lineages by their antagonistic activities on the regulation of neural and mesodermal gene expression (22).

In this study, we aimed at understanding further the relationships between the processes of cell specification and tissue morphogenesis within the PZ, with a particular attention to cellular mechanisms underlying the tightly regulated balance between maintenance of residing posterior progenitors and production of exiting cells that contribute to formation of mesodermal and neural tissues. By analyzing Sox2 and Bra expression in the PZ of the quail embryo, we show that these proteins are expressed with various levels from one cell to another, thus highlighting an important degree of cell-to-cell heterogeneity in this area. Using overexpression and downregulation approaches, we provide evidence that the relative levels of Sox2 and Bra proteins is a key determinant for posterior progenitor choice to stay in place or exit the PZ to join their destination tissues. Time-lapse experiments further revealed that most posterior progenitors are highly migratory without strong directionality. Mathematical modeling and functional experiments then revealed that heterogeneous levels of Sox2 and Bra translate into progenitor specific destiny *via* the control of cell motility: Bra^high^ progenitors display high motility and leave the PZ to join the PSM, Sox2^High^ progenitors are less motile and get integrated into the NT, while progenitors expressing intermediate/equivalent levels of the two proteins tend to remain resident of the PZ.

## Results

### Levels of Sox2 and Bra proteins display high spatial cell-to-cell variability in the PZ

The transcription factors Sox2 and Bra are known to be co-expressed in progenitors of the PZ (19,20). As they are specified from posterior progenitors, neural cells maintain Sox2 expression and downregulate Bra while mesodermal cells downregulate Sox2 and maintain Bra expression. Although Sox2 and Bra are recognized to be key players in driving neural and mesodermal cell fates, the spatial and temporal dynamics of these events remains to be elucidated. As a first step to address this question, we carefully examined expression levels of the two proteins in the PZ of the quail embryo at stage HH10-11. As expected, analyses of immunodetection experiments revealed co-expression of Sox2 and Bra in nuclei of all PZ cells (Fig. 1 A-C) (n=8 embryos). Noticeably, we observed a high heterogeneity in the relative levels of Sox2 and Bra proteins between neighboring PZ cells. We indeed found intermingled cells displaying high Sox2 (Sox2^high^) and low Bra (Bra^low^) levels or, conversely, Bra^high^ and Sox2^low^ levels as well as cells in which both proteins appear to be at equivalent levels. This cellular heterogeneity was very apparent when compared to the adjacent nascent tissues, i.e. the NT and the PSM, where Sox2 and Bra protein levels were found to be very homogenous between neighboring cells (Fig. 1 D-F). Cell-to-cell heterogeneity in posterior progenitor was detected as early as stage HH5-6, a stage corresponding to initial activation of Sox2 and Bra co-expression in the quail embryo (Supplemental Fig. 1). We also observed heterogeneous levels of Sox2 and Bra proteins in PZ cells of chicken embryo, indicating that it is not a specific feature of quail (Supplemental Fig. 2). To infer how Sox2 and Bra protein levels goes from being co-expressed in a heterogeneous manner in the PZ to being expressed homogeneously in the nascent tissues, we analyzed variations of their respective levels in a series of seven volumes (containing around 100 cells in each volume) located in a posterior to anterior path (from the PZ to the maturating tissues), corresponding to putative trajectories of PSM or NT cells (Figure 1G-G’). Data showed that the average expression level of Sox2 increases (+ 2.22 folds, n = 7 embryos) while that of Bra decreases (− 3.81 folds) following the neural path (Figure 1G). On the contrary, along the paraxial mesoderm path, average expression level of Sox2 decreases (− 2.12 folds, n =7 embryos) while Bra level first increases in the posterior PSM (1.14 folds, position 1 to 2) and decreases anteriorly (− 5.06 folds, position 2 to7) (Figure 1 G’). Next, to define whether the cellular heterogeneity found in the PZ depends more on variability of one of the two transcription factors, we quantified protein levels per nuclei of cells populating the PZ. By plotting Sox2 and Bra levels in individual cells, we noticed a broader distribution for Sox2 levels (coefficient of variation of 41.8%) compared to Bra levels (coefficient of variation of 30.75%) (Fig 1 H), indicating that the cell-to-cell heterogeneity in the PZ is preferentially driven by differences in Sox2 levels. To quantify Sox2 and Bra heterogeneity, we calculated the Sox2-to-Bra ratio (Sox2/Bra) for each cell of the PZ as well as for cells of the NT and PSM, and compared these values. Our data showed high divergences between the three tissues and confirmed the high heterogeneity previously observed in PZ cells (Fig 1I). It must however be noticed that these quantitative data revealed a broad range of cell distribution, highlighting, in particular, the presence of cells in the PZ displaying similar Sox2/Bra values as mesodermal or neural cells. We next asked whether the cellular heterogeneity caused by differences in Sox2 and Bra levels is present in the whole volume of the PZ or displays regionalization in this tissue. To address this issue, we analyzed spatial distribution of the Sox2/Bra values on optical transverse sections performed at anterior, mid and posterior positions of the PZ (Figure 1J-J”’). This analysis confirmed the heterogeneity of Sox2/Bra values which are equally represented in the mid area of the PZ (Figure 1J”). Cells with a high ratio level (Sox2^High^ Bra^Low^) were found to be more represented in the most dorso-anterior part of the PZ (Figure 1J’) and cells with a low ratio level (Bra^High^ Sox2^Low^) were found to be more represented in the most posterior part of the PZ (Figure 1J”’). This particular antero-posterior distribution was further confirmed by tissue expression analysis (Supplemental Fig. 3). However, it should be noted that variations of Sox2/Bra values were noticed in all these areas, indicating that the Sox2/Bra-related cell-to-cell heterogeneity is present in the whole PZ.

**Figure 1:**
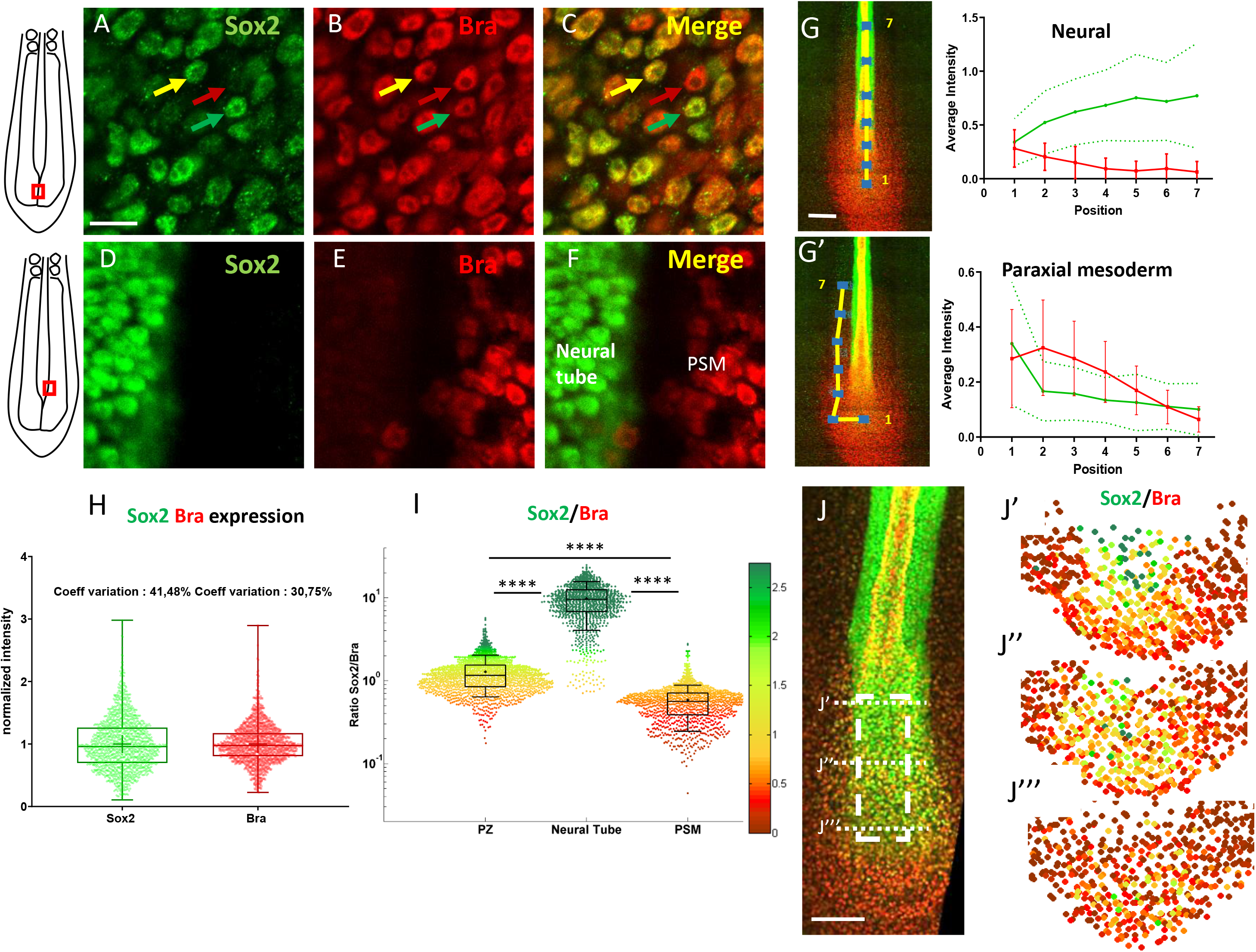
Posterior progenitors co-express Sox2 and Bra with a high degree of cell-to-cell heterogeneity. **A-F** : Immunodetection of Sox2 (green) and Bra (red) analyzed at the cellular scale in the caudal part of stage HH11 quail embryo, either in the PZ (**A-C**) or in the nascent NT and the PSM (**D-F**). Overlay images are presented in **C** and **F**. Note cell-to-cell heterogeneity in Sox2 and Bra levels in the PZ, with neighboring cells expressing higher level of Bra (red arrow), higher level of Sox2 (green arrow) or comparable levels of both proteins (yellow arrow), a feature not apparent in the nascent NT and PSM tissues. **G, G’** : Measurements of Sox2 and Bra levels along putative trajectories of NT (G) and PSM (G’) cells, depicted by yellow lines on whole-mount embryos (left images). Sites of fluorescence measurements are indicated by blue squares (left image), numbered from 1 to 7. **H** : Distribution of normalized cell-to-cell expression of Sox2 and Bra in the PZ (n=8 embryos). Note higher coefficient of variation for Sox2 (41.48%) than for Bra (30.75%). **I** : Cell distribution of Sox2/Bra levels in the PZ (n=9 embryos), the NT (n=7 embryos) and the PSM (n=8 embryos). **J-J”’** : Representation of the Sox2 to Bra ratio (green to red) in digital transversal sections (40 μm) made in the PZ (dashed lines in the double immunodetection image in **J**). Scale bars = 10 μm in **A-F**, 100 μm in **I, L L’.**

Altogether, our data, highlighting significant variability in Sox2 and Bra protein levels within progenitors of the PZ, evidence an unexpected cell-to-cell heterogeneity of this cell population. Noticeably, despite an overall enrichment of Sox2^high^ cells in the dorsal-anterior part of the PZ and Bra^high^ cells in the most posterior part, no clear spatial regionalization of these cells was detected, indicating that the PZ is composed of a complex mixture of cells displaying variable Sox2/Bra levels. This variability is further lost as cells enter the NT or the PSM.

### Relative levels of Sox2 and Bra in PZ cells influence their future tissue distribution

The fact that cell-to-cell heterogeneity caused by differences in the Sox2 and Bra levels is observed in PZ cells but not in PSM and the NT cells was suggestive of a role of these relative protein levels in the decision to leave or not the PZ and to locate in a specific tissue. To test this possibility, we developed functional experiments aimed at increasing or decreasing Sox2 and Bra levels in PZ cells. In the early bird embryo (stage HH4-7), the future posterior progenitors are initially located in anterior epithelial structures: the epiblast and the primitive streak. We thus performed targeted electroporation of progenitors in the anterior primitive streak/epiblast of stage HH5 embryos to transfect expression vectors or morpholinos and further analyzed subsequent distribution of targeted cells, focusing on the PZ, the PSM and the NT (Figure 2). As early as 7hrs after electroporation, we could detect the expected modifications of Sox2 or Bra expression in PZ cells for both overexpression and downregulation experiments (Supplemental Figure 4, 5). We observed a significant decrease in the Sox2/Bra levels by either overexpressing Bra or downregulating Sox2 and a significant increase of this ratio when Sox2 was overexpressed or when Bra was downregulated (Figure 2 A, F). Consistent with previous studies (22), we found that cross-repressive activities of Sox2 and Bra contributed to amplify such ratio’s modifications (Supplemental Figure 5). After transfection of expression vectors or morpholinos, we next let the embryos develop until stage HH10-11 and examined fluorescent cell distribution in the different tissues. For this, we measured the fluorescence intensity of the reporter protein (GFP) in the PZ, the PSM and the NT and calculated the percentage of fluorescence in each tissue. We obtained reproducible data using control expression vector with less than 20% of the fluorescent signal found in the PZ (16.78% ±2.83), a little more than 20% in the PSM (22.64 % ± 3.30) and about 60% in the NT (60.57% ± 4.39) (Figure 2 B, E). We next found that overexpression of Bra leads to a marked reduction of the fluorescent signal in the PZ (1.17% ± 0.57) and to an increased signal in the PSM (33.04% ± 4.06) but has no effect on the NT signal (Figure 2 B, C, E). Elevating Bra levels is thus sufficient to trigger cell exit from the PZ and to drive cells to join the PSM. However, this is not sufficient to impede PZ cell contribution to form the NT. Similarly, we found that overexpression of Sox2 drives exit of the cells from the PZ (1.16% ± 0.67) favoring their localization in the NT (75.40% ± 4.57) without significantly affecting proportions of cells in the PSM (Figure 2 B, D, E). Spatial distribution of the fluorescent signals obtained using control morpholinos appeared very similar to that observed using the control expression vector (18.93% ± 3.06, 22.68 ± 4.09 and 58.38% ± 3.63 for the PZ, the PSM and the NT, respectively) (Figure 2 G, J). We found that downregulation of Bra leads to exit of cells from the PZ (4.16% ± 1.57) and favors cell localization in the NT (88.23%, ± 2.04) at the expense of the PSM (7.59% ± 1.81) (Figure 2 G, H, J). Similarly, Sox2 downregulation triggers cell exit from the PZ (7.50% ± 2.35) and, as expected, leads to higher contribution of cells to the PSM (34.59% ± 4.79) but this does not occur at the expense of cell contribution to the NT (Figure 2 G, I, J).

**Figure 2:**
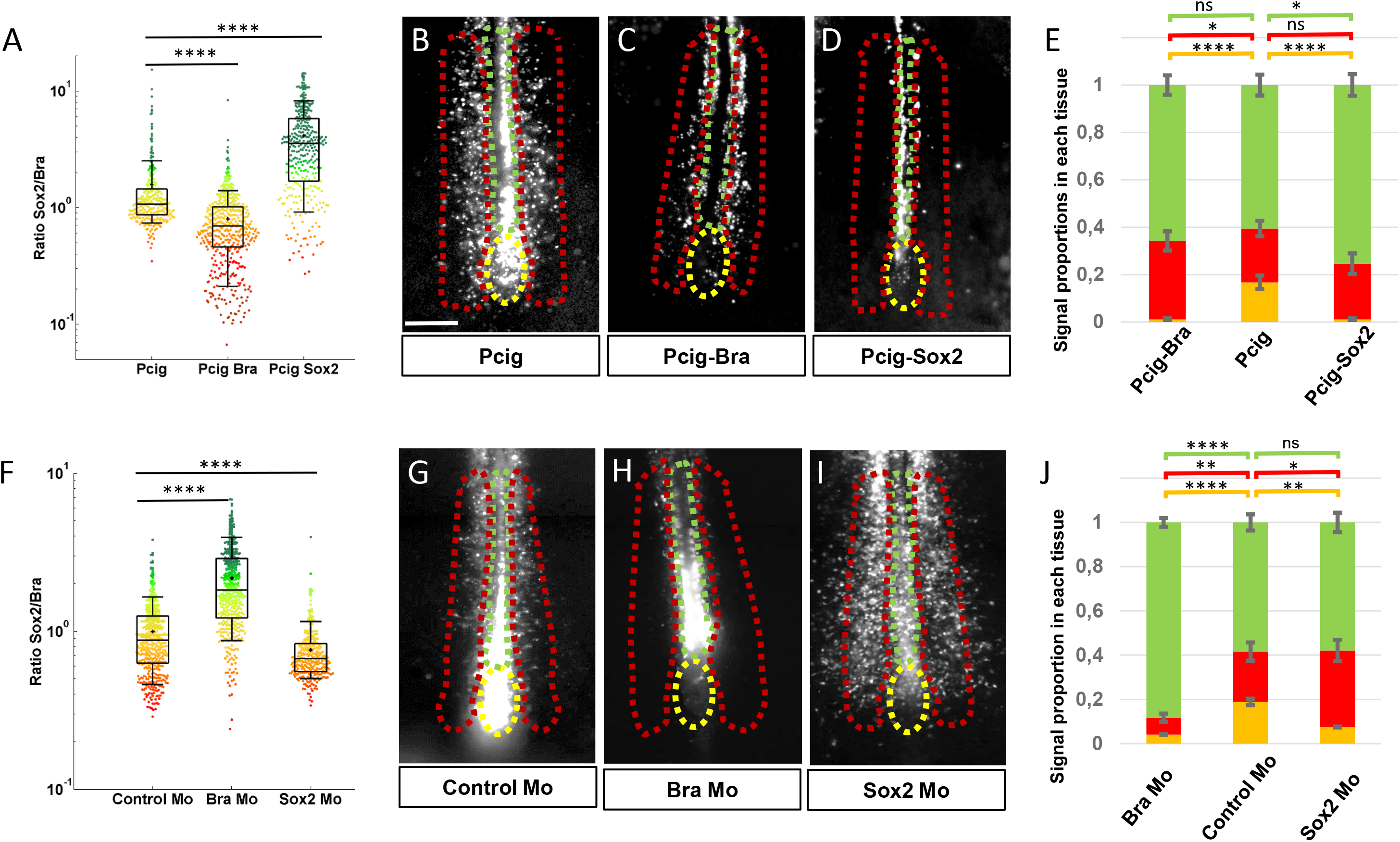
Sox2 and Bra levels are critical for progenitor maintenance and tissue distribution. **A, F:** Sox2 to Bra ratios calculated following Bra and Sox2 double immunodetection in the PZ performed 7 hours after electroporation. Values were normalized to the average ratio of non-transfected cells of the same region. **A:** Sox2/Bra values in cells transfected with Bra (Pcig-Bra) and Sox2 (Pcig-Sox2) expression vectors compared to cells transfected with the empty vector (Pcig). **B:** Sox2/Bra values in cells transfected with morpholinos directed against Bra (Bra Mo) or Sox2 (Sox2 Mo) compared to cells transfected with a control Mo. Ratios were calculated on the basis of 286 to 590 cells and 3 to 5 embryos per condition. **B-D, G-I:** Ventral views of embryos collected 20 hours after electroporation showing the GFP signals (white). The PZ, the PSM and the NT are delineated by yellow, red and green dash lines, respectively. Expression vectors or morpholinos used are indicated below each picture. Scale bar = 100 μm. **E, L:** Staked histograms displaying the proportion of cells in the PZ (yellow), the PSM (red) and the NT (green). For each experimental condition, proportion of cells in a given tissue was compared to the same tissue of control embryos by unpaired Student test (n= 27 embryos for Pcig Bra, n= 21 embryos control for Pcig, n=23 embryos for Pcig Sox2; n=28 embryos for Bra Mo, n=27 embryos for Control Mo and n=28 embryos for Sox2 Mo). Errors bars represent the SEM.

These data, showing that changing the Sox2 to Bra ratio, tending either towards higher or lower values, is sufficient to trigger cell exit from the PZ, evidence that the relative levels of Sox2 and Bra proteins are key determinant of PZ cell choice to stay in the PZ or exit this area to enter more mature tissues. Our data also point to the critical influence of the relative levels of Sox2 and Bra in controlling the final destination of cells exiting the PZ, with Sox2^high^ (Bra^low^) cells and Bra^high^ (Sox2^low^) cells preferentially integrating the NT and the PSM, respectively. Intriguingly, we also observed that the preferential cell colonization of one given tissue is not always clearly associated with depletion of cells in the other (see discussion).

### PZ cells are highly motile without strong directionality

To better characterize the movements of posterior progenitors, either staying resident to the PZ or exiting this area, we examined their behaviors using live cell imaging. We electroporated stage HH5 quail embryos with a vector encoding for nuclear GFP and performed time-lapse imaging experiments from stage HH8 to stage HH12. At these stages (stage HH8 and onward), posterior progenitors are no longer located in the dorsal epithelium but rather within a dense and internal mesenchymal structure that prefigure the embryonic tailbud (12,23). In order to compare migration properties between tissues, we focused on the PZ and on the PSM and the posterior NT (Figure 3A, Supplemental Movie 1). Because the three tissues have a global movement directed posteriorly due to the embryonic elongation, we generated two types of cellular tracking: the raw movement, in which the last-formed somite is set as a reference point, and the “corrected” movement, in which the cellular movements are analyzed in reference to the region of interest (Figure 3B). Tracking cell movements allowed for quantification of motility distribution, directionality of migration and time-averaged mean squared displacement (Figure 3C-E) (n=7 embryos). First, we noticed that the average raw motility of PZ cells is higher than that of PSM or NT cells (Figure 3C, top panel). Raw directionality was also found more pronounced for PZ cells in the posterior direction compared to PSM or NT cells (Figure 3D, upper panel). These results thus confirm that the PZ is moving faster in a posterior direction than surrounding tissues, as previously measured using transgenic quails embryos (24). Analysis of local (corrected) motility reveals that PZ cells move in average as fast as PSM cells and significantly faster than NT cells (Figure 3C, bottom panel). The distribution of individual corrected PZ cell motilities is however different from the ones of PSM cells as analysis in the PZ showed slower moving cells (PZ corrected motility violin plot in Figure 3C is larger for slow values than the PSM counterpart and Supplemental Figure 6), indicating motile behavior of PZ cells is more heterogeneous than that of PSM cells. As previously reported (25), we found that, after tissue correction, the directionality of PSM cell motion was mostly non-directional with, however, a slight tendency towards anterior direction which is expected due to the posterior elongation movements of the reference tissue (Figure 3D, red plot in the lower panel). The distribution of corrected angles of PZ cell motilities was also found globally non-directed, with however a slight tendency towards anterior direction, to some extent more pronounced than for PSM cells, suggesting that our method is able to detect trajectories of cells exiting the PZ to integrate the NT or the PSM (Figure 3D, yellow plot lower panel). Examination of individual cell tracks further confirmed extensive non-directional local migration and neighbor exchanges within the PZ (Supplemental Movie 2). As PZ cell movement was found being mostly non-directional, we next looked at their diffusive motion by plotting their mean squared displacements (MSD), measured in each tissue over time, as it has been previously done for PSM cells (25). This analysis showed that the MSD of posterior progenitors is linear after tissue subtraction, as intense as the MSD of PSM cells and significantly higher than that of NT cells, thus demonstrating the diffusive nature of PZ cell movements (Fig. 3E).

**Figure 3:**
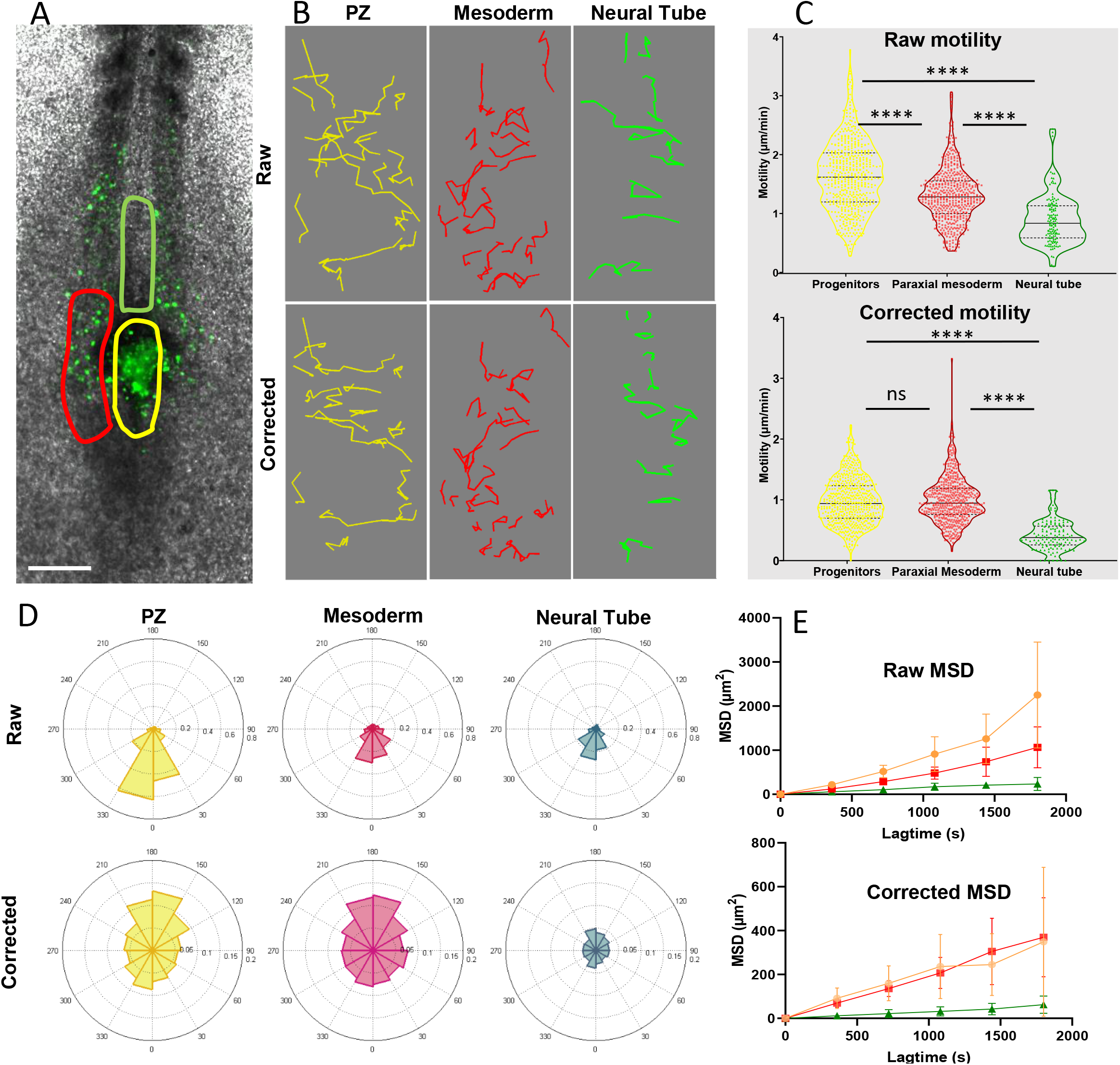
Progenitors display high motility without strong directionality. **A.** Representative image of a H2B-GFP electroporated quail embryo (ventral view) analyzed by live imaging analysis. Transfected cells are detected by the GFP signal (green) and the PZ, the nascent PSM and the NT are delineated by yellow, red and green lines, respectively. **B**: Example of cell trajectories before (raw) and after tissue motion subtraction (corrected). **C**. Distribution of the raw (top) and corrected (bottom) cell motilities computed in the different regions. **D**. Directionality of motion assessed by the distribution of angles weighed by the velocity for the different regions, before and after tissue subtraction. **E.** Assessment of diffusion by analysis of the mean squared displacement in function of time for the different regions (n=7 embryos, 538 cell trajectories analyzed in the PZ, 496 in the PSM, 128 in the NT). Scale bar = 100 μm.

Together, these data evidenced that, in the referential of the progenitor region, PZ cell migration is diffusive/without displaying strong directionality (except a slight anterior tendency), with an average motility that is comparable to that of PSM cells and significantly higher than that of NT cells. The motility of individual PZ cells however appears heterogeneous with some cells exhibiting high motile behavior, as do PSM cells, and others displaying low motility comparable to that of NT cells.

### Modeling spatial cellular heterogeneity and tissue morphogenesis

To go further in understanding how variation in Sox2 and Bra protein levels between posterior progenitors might control their choice of residing or exiting the PZ, we designed an agent based mathematical model (Figure 4). In this model, we considered biological data showing spatial heterogeneity of Sox2/Bra levels within the population of PZ cells. Furthermore, based on our data showing that variable motile behaviors exist within the population of PZ cells, resembling either that of PSM or NT cells, we assumed that PZ cell motility is directly driven by the Sox2/Bra value. Working hypothesis was that Bra promotes non-directional motility while, conversely, Sox2 inhibits it. We chose a time window corresponding to stages HH8-HH12, a developmental period when the NT, the PSM and the PZ have already been formed. To consider the physical boundaries existing between these tissues, we integrated a non-mixing property between cell types. As there is few dorso-ventral tissue deformations during the selected time window (21), we designed a 2D model (X,Y). We implemented cell numbers, cell proliferation and tissue shape to be as close as possible to biological measurements (21) (Supplementary methods). The model is set up in a way that PZ cells express random and dynamic Sox2/Bra levels with a defined probability to switch into a Bra ^high^(Sox2^low^) state (PSM state) or into a Sox2 ^high^(Bra ^low^) state (NT state). The motility is directly controlled by the Sox2/Bra ratio: Bra ^high^(Sox2 ^low^) cells display high motility, Sox2 ^high^(Bra ^low^) cells display low motility and undetermined progenitors display intermediate motility (Figure 4A). We first verified that our mathematical model recapitulates the basic properties of the biological system: in particular, it allows for recreating spatial heterogeneity of the relative Sox2/Bra levels in cells of the PZ (Figure 4B) and reproduces general trends in tissue motility and non-directionality (Figure 4C, D). Simulations of our model also showed that the relative cell numbers (taking into account of proliferation) evolve as expected with a stable number of PZ cells and an increased number of NT and PSM cells (Figure 4E). We next explored the ability of the model to reproduce maintenance of residing posterior progenitors while the NT and PSM extend towards the posterior pole during the elongation process. Looking at different time points of the simulation process, we indeed observed that the PZ is maintained posteriorly during the elongation process (Figure 4F, Supplemental Movie 3). This mathematical model thus reproduces with success the main properties of the biological system. Noticeably, simulations also showed dynamic deformation of the PZ, which adopts asymmetric shapes and then gets back to a symmetric shape (Supplemental Movie 3), thus highlighting self-corrective properties of the system. To check if spatial heterogeneity of Sox2/Bra levels can self-organize in our model, we made a simulation in which each posterior progenitor starts with equivalent levels of Sox2 (50%) and Bra (50%). We observed that spatial heterogeneity emerges almost immediately and persists all along the simulation, suggesting that this feature self-organizes independently of the initial levels of Sox2/Bra values (Supplemental Movie 4). To challenge further this model and test if it can recapitulate the experimental results we obtained by over-expressing or down-regulating Sox2 and Bra, we explored the consequences of numerically deregulating the Sox2/Bra values on tissues and cell behaviors. As a result, Bra^High^ values increase PZ cell motility in the model (Figure 4G), lead to generation of a higher number of PSM cells (Figure 4H), a depletion of cells in the PZ and a shorter NT (Figure 4H, J; Supplemental Movie 5). At the opposite, Sox2^High^ values lead to a reduced PZ cell motility (Figure 4G), a depletion of PZ cells and an increased number of NT cells and width (Figure 4I, J; Supplemental Movie 6). Results of numerical simulations therefore agree with experimental data showing that Sox2/Bra levels control the balance between maintenance of progenitors in the PZ and continuous distribution of cells into the NT and the PSM. Together, data from our mathematical simulation strongly support the view that the distinct progenitor behaviors might be guided by Sox2/Bra-dependent heterogeneous cell motility.

**Figure 4:**
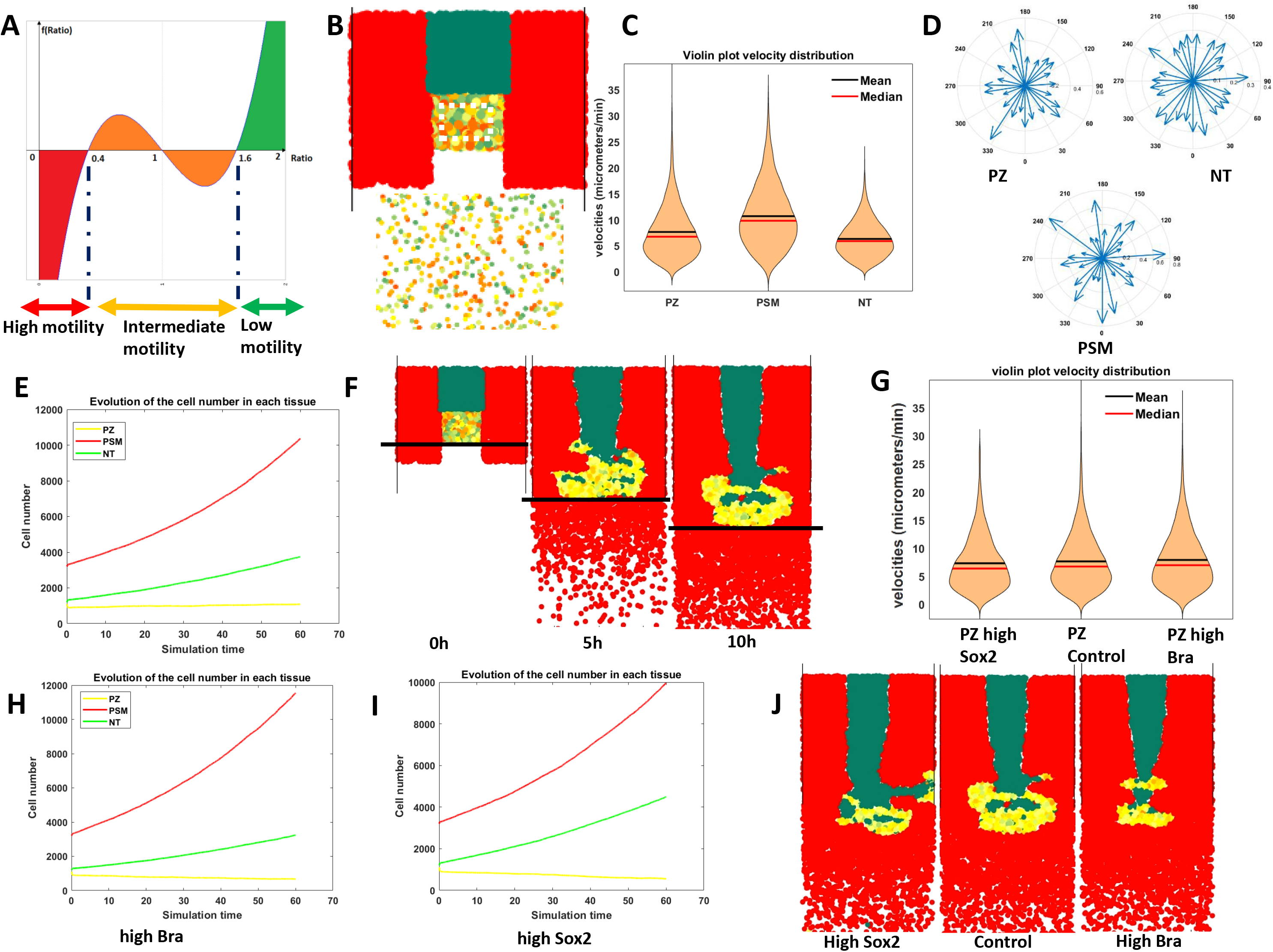
Mathematical modeling of posterior progenitor behavior downstream of heterogeneous expression of Sox2 and Bra. **A**: Graphical representation of the mathematical function defining the Sox2 to Bra ratio dynamics. In each progenitor cell, the Sox2/Bra value oscillates randomly from 0.4 to 1.6 and noise in the system ensures that some cells pass below 0.4 to be specified into PSM cells (red) while some cells pass above 1.6 to become NT cells (green). Low ratios (below 0.4) confer high-motility, high ratios (above 1.6) inhibit motility, ratios between 0.4 and 1.6 confers intermediate level of motility. **B:** Progenitor region showing the spatial heterogeneity of Sox2/Bra levels with a close up on the PZ on the bottom panel. **C**: Distribution of cell motilities within the PZ, the PSM and the neural tube (NT). **D**: Directionality of migration in the three tissues. **E**: Evolution in the number of each cell type over time. **F:** Simulation of PSM (red), NT (green) and PZ (yellow) evolution at the time points of 5 and 10 hours. Note that the different tissues are preformed at the initial time (0h) and that tissues elongate over time (compare positions of the black bars). **G-J**: Effects of numerical deregulations of the Sox2/Bra values, i.e. imposing low Sox2/Bra (high Bra) or high Sox2/Bra (high Sox2) values, on the distribution of cell motility (**G**), cell numbers (**H, I**) and on tissue evolution at 5 hours (**J**). Each parameter was analyzed in comparison with data obtained using the random model (control).

### The Sox2 to Bra ratio controls motility of PZ cells

To test the prediction that the relative levels of Sox2 and Bra direct their choice of staying in place or exit the PZ and contribute to NT or PSM formation, we developed functional experiments *in vivo*. Sox2 and Bra were either over-expressed or down-regulated in PZ cells and the behaviors of posterior progenitors were followed by time-lapse imaging (Figure 5 A-F). We first monitored raw cell motilities (Supplemental Figure 7) and conducted subtraction of the tissue motion to gain insight into local motility and directionality (Figure 5). We found that Bra-overexpressing PZ cells display higher motility without significant differences in directionality when compared to control cells. By contrast, when PZ cells overexpress Sox2, we detected a significant reduction of their motility accompanied by an anterior bias in angle distribution compared to control cells (Figure 5 B, C, D, and Supplemental Movie 7). We found that Bra down-regulation leads to similar significant reduction of cell motility, as well as a change in directionality towards the anterior direction (Figure 5 B, E, F). Conversely, Sox2 down-regulation did not result in significant effect on average cell motility or directionality, even though a tendency towards a slight increase in motility was noticed (Figure 5 B, E, F; Supplemental Movie 8).

These data, showing that changing the respective levels of Sox2 and Bra is sufficient to modulate PZ cell motility/migration properties, highlight a key role for these transcription factors in controlling PZ cells movements with Sox2 and Bra inhibiting and promoting cell motility, respectively. When cells have high Sox2/Bra levels they migrate less and are left behind the PZ to be integrated into the NT. When cells have a low Sox2/Bra ratio they tend to migrate more, mostly in a diffusive manner, explaining how they leave the PZ to be integrated in the surrounding PSM tissues.

**Figure 5:**
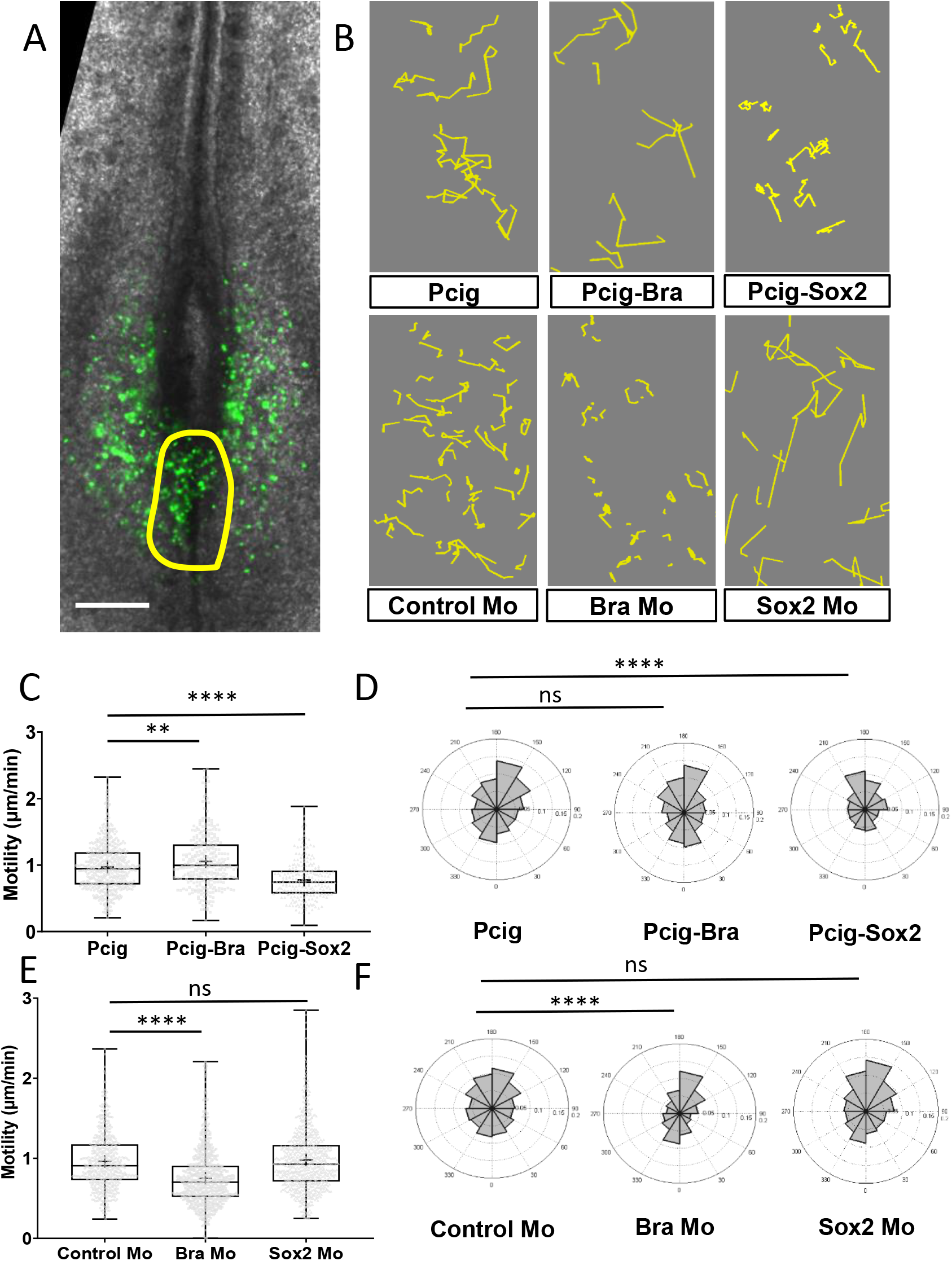
Sox2 and Bra deregulation affect progenitor motility. **A**: Representative image of a Pcig electroporated quail embryo (ventral view) used to perform progenitor tracking and motility analysis. Transfected cells are detected by the GFP signal (green) and the PZ is delineated by the yellow line. **B**: Examples of cell tracks after correction in embryos electroporated with expression vectors or morpholinos indicated on each panel **. C, E:** Distribution of PZ cell motilities after tissue motion subtraction in gain of function (**C**) and in down-regulation (**E**) experiments. **D, F:** Directionality of cell motion after tissue motion subtraction assessed by the distribution of angles in gain of function (**D**) and in downregulation (**F**) experiments (n=7 embryos and 541 trajectories for Pcig, n= 5 embryos and 307 trajectories for Pcig-Bra, n= 5 embryos and 234 trajectories for Pcig-Sox2; n=5 embryos and 590 trajectories for Control Mo, n=7 embryos and 753 trajectories for Bra Mo, n=5 Embryos and 874 trajectories for Sox2 Mo). Scale bar = 100 μm.

### Modeling the importance of spatial cell-to cell heterogeneity in morphogenesis

Results of our mathematical model argue in favor of random spatial distribution of PZ cells being important to establish the proper balance between their two choices of staying in place or exit the PZ. To validate this model, we decided to compare it to second model in which cell-to-cell heterogeneity is spatially organized. As shown above (Figures 1J and Supplemental Figure 3), cells harboring either Sox2^High^ or Bra^high^ levels were found intermingled within the PZ without a clear pattern except in the dorsal part of the PZ where Sox2^High^ and Bra^high^ cells are organized following two opposite gradients: an antero-posterior decreasing gradient of Sox2 and an antero-posterior increasing gradient of Bra.

According to this observation, we chose to impose Sox2 and Bra levels to be distributed in such opposite gradients in a second model (Figure 6A). This new version of the model, named the gradient model, has been set up to reproduce similar dynamics in cell specification (Figure 6B) and relationships between Sox2/Bra values and motility as in the random model (Supplemental methods). We noticed that cell motility in the gradient model was globally comparable to that of the random model (Figure 6C, 4C) and was mostly non-directional (Supplemental Figure 8). We also observed maintenance of PZ cells caudally and elongation of the different tissues (Figure 6D, Supplemental Movie 9), suggesting that a gradient in Sox2 and Bra expression can also insure proper progenitor maintenance and cell distribution in the NT and PSM. Despite the observed similarities, we however noticed several crucial differences between the two models. Indeed, the speed of elongation was less important in the graded simulation (0.9 a.u.) versus the spatially heterogeneous (random) simulation (1.9 a.u.) (Figure 6E). To test if posterior movements of resident progenitors differ between the two models, we tracked virtual cells that stay in the PZ and calculated the distances they travelled in Y direction. This analysis showed that resident progenitors move more posteriorly in the random (mean distance of 3.32 a.u.) versus gradient model (mean distance of 3.14 a.u.), suggesting that random distribution of Sox2/Bra ratio is more efficient in keeping progenitors in their posteriorly moving niche (the PZ) than graded distribution (Figure 6F). Furthermore, we also noticed fewer transient deformations and self-corrective behavior of the PZ tissue in the gradient simulation when compared to the random simulation (Supplemental Movie 10). By analyzing the PZ shapes between the beginning and the end of the simulation, we found that it was much more conserved in the random model compared to the gradient model. Indeed, it became larger (medio-lateral) and shorter (antero-posterior), showing a conservation of proportions (length/width) of only 37% for the gradient model versus 74% for the random model (Figure 6G). Finally, to test if the changes we observed at the tissue level could be due to changes in the diffusivity of cellular migration, we plotted the MSD through time for both the random and gradient models. We found that the MSD of the random model is higher than that of the gradient model (Figure 6H), suggesting that the spatial heterogeneity in expression of Sox2 and Bra is enhancing the diffusive behavior of PZ cells. This higher diffusivity can therefore bring more tissue fluidity to remodel it more efficiently and to maintain its shape over long time scales. Together, modeling data indicate that random distribution of cell-to-cell heterogeneity is more efficient compared to an organized cell distribution in promoting posterior movements of progenitors, tissue fluidity and PZ remodeling.

**Figure 6:**
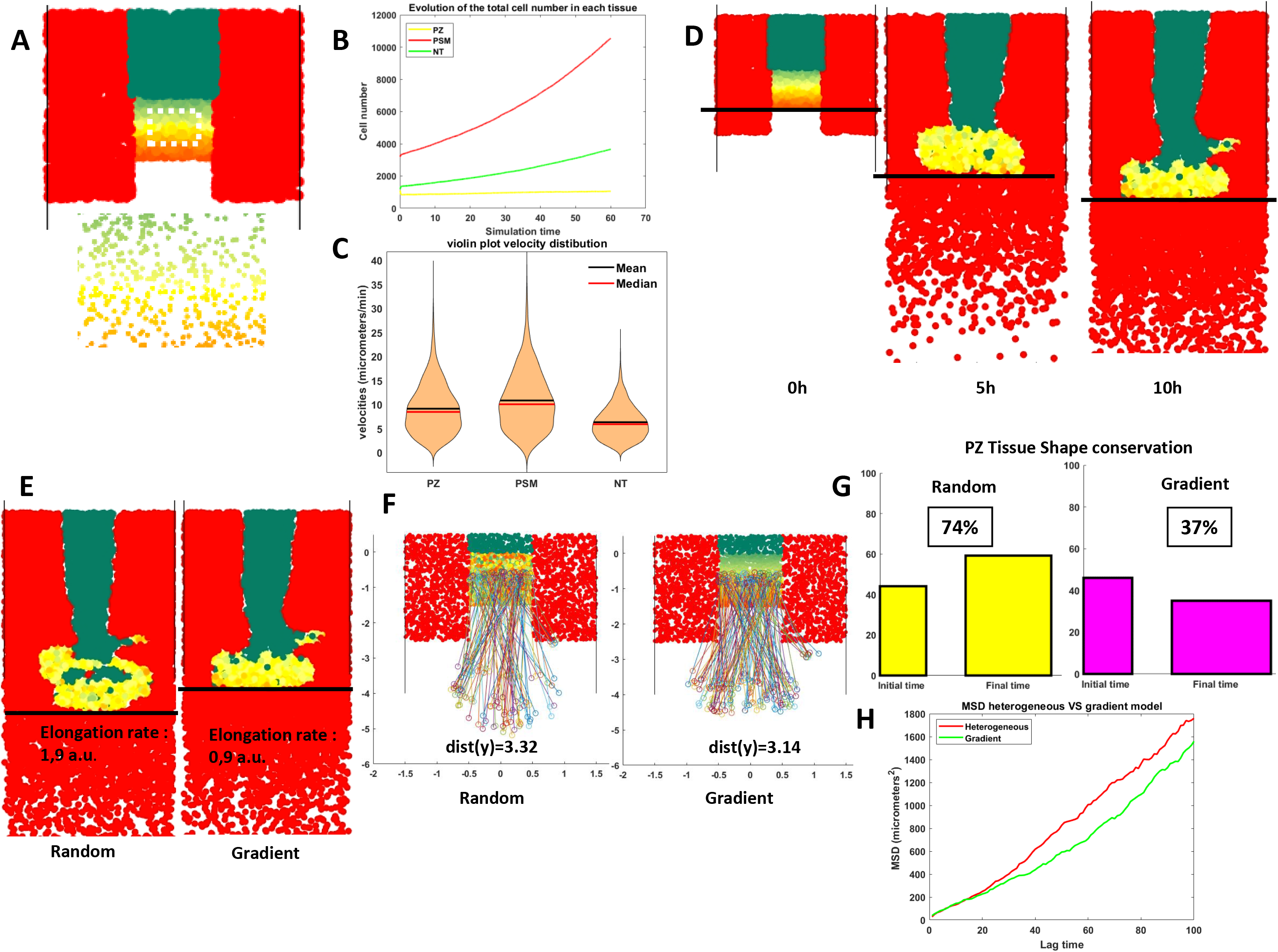
Comparison of spatial heterogeneity and gradient models. **A:** Initial condition of the gradient model and close up of the PZ showing the spatial organization of Sox2/Bra levels, following opposite gradients (bottom panel). **B**: Time evolution of cell numbers for each cell type according to the gradient model. **C**: Distribution of cell motilities within the PZ, the PSM and the NT according to the gradient model. **D** Evolution of the gradient model through the different time points of 0, 5 and 10 hours. **E**: Comparison of the gradient and heterogeneous simulations at 10 hours. Note differences in the elongation rates. **F**: Y displacement of resident progenitors located at the center (along the anteroposterior axis) of the PZ for each model. Values below the graphs are averages of Y displacements. **G:** Initial (left) and final (right) shapes of the PZ according to the heterogeneous (yellow) and the gradient (pink) models. Conservation or proportions (length/width) are noted in percentage (100% would correspond to an unchanged shape). **H**: MSD calculated for progenitors in the heterogeneous (red) and gradient (green) models.

## Discussion

In the present work, using experimental and theoretical approaches, we bring evidence that the spatial cell-to-cell heterogeneity created in the PZ by variations of the Sox2 to Bra ratio is critical to regulate progenitor destiny through the control of their motile behavior. Our data support a model in which high levels of Sox2 give cells low motile properties and make them integrate the NT while high levels of Bra rather give them high motile and diffusive properties that push them to exit the PZ and integrate the PSM. Located in-between these low and high motile/diffusive cells, are progenitors co-expressing Sox2 and Bra at comparable levels, that are moving with an intermediate speed, and that remain resident of the PZ as this tissue is moving posteriorly. This study, by unraveling that specification and morphogenesis are coupled during embryonic axis elongation, shed a new light on how cell-to-cell heterogeneity ensures robustness in morphogenesis.

In this work, we show that Sox2 and Bra proteins are co-expressed in PZ cells of quail embryos. This co-expression is a conserved property of vertebrate embryos since it has been previously reported in chick, zebrafish, mouse and human embryos (19–21). Interestingly, it has also been noticed that Sox2 and Bra are expressed at different levels in the PZ, following, in particular, opposite expression gradients with anterior cells expressing high levels of Sox2 which decays posteriorly and posterior cells expressing high levels of Bra which decays in the opposite direction (20,26). Even though our data showed that such a pattern is apparent in the dorsal part of the quail PZ, an important finding of our work is that cells with variable levels of Sox2 and Bra are found intermingled in all areas of the PZ. How this cell-to-cell heterogeneity is established and further maintained over the elongation process remains an open question. Graded activity of signaling pathways such as Wnt, FGF and RA, all known to regulate Bra and Sox2 expression (27–31), together with dynamic cross-repressive activities of Sox2 and Bra (our present data and (22)) and cellular movements are likely to contribute to create and maintain such a heterogeneity.

The antagonistic interaction between Sox2 and Bra has furthermore been proposed to determine fate decision of posterior progenitors (22). The different levels of Sox2 and Bra expression we evidenced between posterior progenitors is thus indicative of the presence of mixed cell populations in the PZ harboring different specification states: Bra^High^ progenitors being engaged towards the mesodermal fate, Sox2 ^High^ towards the neural fate while progenitors with comparable levels of the two proteins being situated in between these two states. In agreement, we found that forced expression of Sox2 and down-regulation of Bra favor integration of posterior progenitors into the NT while forced expression of Bra and downregulation of Sox2 favor their distribution to the PSM. However and intriguingly, these results could not explain why, in gain and loss-of-function experiments, preferential distribution of electroporated cells into the NT is not always paralleled by a decrease in their participation to PSM formation (or inversely) (Figure 2E, J). A possible explanation is that a progenitor which is already engaged toward a given fate is no longer competent to switch its fate and, thereby, to change its tissue destination. As mentioned above, we found the different types of progenitors intermingled in all areas of the PZ. Single cell sequencing studies have revealed the molecular signatures of the different progenitor states; however, due to technical limitations these studies could not reveal their exact locations within the posterior region. Fate maps studies around stage HH4-5 have shown that the distribution of progenitors along the antero-posterior axis of the epiblast/streak is translated in the distribution of their descendants along the medio-lateral axis in formed tissues of older embryos (NT, PSM, lateral plate)(4,6). In this perspective, anterior cells, which are expressing high levels of Sox2, give rise to neural cells and more posterior cells, expressing high levels of Bra, give rise to PSM (and eventually to lateral mesoderm for cells located even more caudally). The fact that we found Sox2 and Bra heterogeneously expressed within the PZ is rather suggestive of a more complex picture where position in the progenitor region does not systematically prefigure final tissue destination. Following this scenario neighboring progenitors could give rise to progeny in different tissues, an observation that is consistent with prospective maps of the PZ in which a small number of labelled cells participate to distinct tissues (3–6).

Analysis of our time-lapse experiments show that most PZ cells are highly mobile, and that this motility is mainly non-directional. Overexpression of Sox2 or down regulation of Bra strongly inhibits cell motility in the PZ leading to an anterior bias in the direction of progenitor movements. At the opposite, overexpression of Bra and, to some extends downregulation of Sox2, favor a slight increase in PZ cell motility. Similarly, cells located axially in the zebrafish tailbud have been shown to display highly disordered motility, suggesting conservation of the role of high and non-directional progenitor’s motility between vertebrate species (32,33). The fact that posterior progenitors often exchange neighbors offers an explanation on how the spatial heterogeneity of posterior progenitor is sorted out to form PSM and NT. Indeed, thanks to their highly migratory properties, Bra^High^ cells could make their way to the surrounding PSM by moving in between other cells including Sox2 ^High^ cells that are less motile. It has been shown that Brachyury has a role in cell migration (17,18). In particular, mouse cells that have a mutation in the Brachyury gene have lower migration speed than wild-type cells when isolated and cultured, explaining part of the mouse embryonic axis truncation phenotype (34). Although a role for Sox2 in the control of progenitor cell migration has, to our knowledge, not previously been reported, recent works have demonstrated that a rise of Sox2 expression promotes the transition of posterior progenitors to NT during chick embryo secondary neurulation (35) and that turning off Sox2 is necessary for NMP to enter the mesoderm in zebrafish embryo (36). In addition, it has also been observed by time-lapse analysis that the zone between the PZ and the NT does not display excessive migration or neighbor exchanges (37). Together, these data confirm the hypothesis that Sox2^High^ cells could be passively laid down as the PZ move posteriorly. In our experiments, while a clear inhibition of cell motility can be obtained by Bra downregulation and Sox2 overexpression, only a subtle enhancement of cell motility was obtained downregulating Sox2 and overexpressing Bra. These differences can be explained by the fact that posterior progenitor’s Sox2/Bra ratios and motilities are much more similar to ratios and motilities of PSM cells than NT cells (Figure 1,3). Biasing progenitors with mesenchymal properties towards a neural state is therefore much more likely to give a difference in motility than a change towards another mesodermal state. In lines with this explanation is the fact that during the course of axis elongation posterior progenitors undergo an epithelial-mesenchymal transition before reaching their full potential to give rise to progeny in the NT and the PSM (12,31,38). Therefore, it is likely that even though Sox2/Bra heterogeneity is present since stage HH5, regulation of progenitor destiny by cellular motility is mostly active after stage HH8 when the progenitors have become mesenchymal with cellular properties that are closer to PSM cells. Indeed, we observed that PZ cell motilities were higher if analyzed between stages HH8 to HH12 compared to the earlier stages HH5 and HH8 (data not shown). Interestingly, the posterior global movement of the PZ region seems constant all along those different stages. Several works, realized in bird embryo, have indicated that physical constrains exerted by neighboring tissues, in particular the PSM, promote the posterior movement of the PZ (24,25,39,40). It is therefore likely that the posterior movement of PZ cells is the result of both local re-arrangements and external forces acting on the whole region.

Based on simulations we got from the two mathematical models, we propose that both a spatially random and a patterned heterogeneity in Sox2 and Bra expression are able to maintain progenitors caudally and to guide their progeny in the NT and the PSM. In our biological system, we observe a superposition of spatially random and a patterned distribution of Sox2/Bra levels. All these data thus support the view that both systems are simultaneously at work in the embryo. However, little is known about the role of spatial cell-to-cell heterogeneity during morphogenesis. Here, we propose that the spatially random pattern allows more posterior movements, cell rearrangements and tissue fluidity in the progenitor zone. This fluid-like state and the opposite solid-like state of the anterior PSM tissue have been shown to be key for zebrafish embryo axis elongation (32,41). In addition, the more efficient self-correction observed in the random model is also supportive of spatial cell-to-cell heterogeneity in the PZ providing plasticity to the system. Several studies have shown that this particular region of the embryo is able to re-generate after partial ablation (42,43). Spatial cell-to-cell heterogeneity, which allows easier re-organization of remaining cells than graded cell pattern, thus appears to be an enabling factor for self-correction. This is even the more so considering that if gradients of Sox2 and Bra are controlled by secreted signals, tissue ablation could be more detrimental to the diffusion of these signals (and re-patterning) than auto-organization of cell-to-cell heterogeneity. Spatial heterogeneity in gene and protein expression is a common trait of living systems and have been observed in many contexts including cancer cells (44,45). The link between cellular spatial heterogeneity and the robustness of morphogenetic processes that we describe here can therefore be relevant beyond the scope of developmental biology.

## Material and Methods

### Quail Embryos and cultures

Fertilized eggs of quail (Coturnix japonica), obtained from commercial sources, were incubated at 38°C at constant humidity and embryos were harvested at the desired stage of development. The early development of quail being comparable to chicken, embryonic stages were defined using quail (46) or chicken embryo development tables (47). Embryos were grown *ex ovo* using the EC (early chick) technique (48) for 6 to 20 hours at 39°C in a humid atmosphere.

### Expression vectors and morpholinos

cBra full length cDNA was cloned by PCR using the following primers (5’-ACCATGGGCTCCCCGGAG-3’; 5’-CTACGCAAAGCAGTGCAGGTGC-3’) into Pcig (49). cSox2 was cloned from Pccags-cSox2 (gift from Daniela Roellig (50)) using EcoRV/XbaI into Pcig to obtain Pcig-cSox2. Fluorescein-coupled morpholinos (Mo) were synthesized by Gene Tools. The nucleotide sequences of the morpholinos were designed to target the translation initiation site of quail Bra (5’-AAATCCCCCCCCCCTTCCCCGAG-3’) and Sox2 (5’-GTACATTCAAACTACTTTTGCCTGG-3’) mRNAs. The Mo (5’-CCTCTTACCTCAGTTACAATTTATA-3’) directed against the transcript of β-human globin was used as control.

### Electroporation

We collected stage 4-6 quail embryos. The solution containing the morpholinos (1mM) and pCIG empty (1-2μg/μL) as a carrier or the DNA solution containing expression vectors Pcig, pCIG-Bra or pCIG-Sox2 (2-5μg/μL) were microinjected between the vitelline membrane and the epiblast at the anterior region of the primitive streak (51). The electrodes were positioned on either side of the embryo and five pulses of 5.2V, with a duration of 50ms, were carried out at a time interval of 200ms. The embryos were screened for fluorescence and morphology and kept in culture for up to 24 hours. To observe the distribution of fluorescence in electroporated tissues, embryos were cultured overnight and fixed before being mounted, ventral side up.

### Immunodetection

For immunodetection, embryos of stages HH 9 to HH11 were fixed for 2 hours at room temperature in formaldehyde 4% in PBS. Blocking and permeabilization were achieved by incubating the embryos in a solution containing Triton X-100 (0.5%) and donkey serum (1%) diluted in PBS for 2 hours. The embryos were then incubated with primary antibodies to Sox2 (1/ 5000, EMD Millipore, ab5603) and Bra (1/500, R&D Systems, AF2085) overnight at 4°C under agitation. After washes, the embryos were incubated with secondary antibodies coupled with AlexaFluor555, AlexaFluor488 (1/1000, ThermoFisherScientific) and with DAPI (4’,6-diamidino-2-phenylindole, 1/1000, ThermoFisherScientific D1306) overnight at 4°C under agitation.

### Image acquisition and processing

Image acquisition for immunodetection was performed using a Zeiss 710 laser confocal microscope (20x and 40x objectives). Quantification of Sox2 and Bra levels in 3D was made with Fiji or with the spot function (DAPI Staining) of Imaris. Immunodetection signals were normalized to DAPI signal to consider loss in intensity due to depth of the tissue. Immunodetection signals and ratios were calculated and plotted using Matlab. Quantification of protein levels in gain and loss of function experiments was performed 7 hours after electroporation by analysing immunodetection signal levels within GFP positive progenitors and by normalizing to endogenous expression measured in non-electroporated cells. Fluorescence distribution in tissues was acquired on a wide field microscope Axio-imager type 2 (Colibiri 8 multi-diode light source, 10X objective). Images of electroporated embryos were processed with the Zen software that allows the assembly of the different parts of the mosaic (“Stitch” function) and were then processed with the “Stack focuser” plugin of the Image J software. The different tissues were delineated on ImageJ with the hands-free selection tool and the images were then binarized using the threshold tool. The total fluorescence intensity emitted by cells transfected with the different constructs was measured and the sum of the positive pixels for the different tissues was calculated. The percentage of fluorescence distribution in the different tissues was then calculated.

### Live imaging and cell tracking

Live Imaging was done using Zeiss Axio-imager type 2 (10X objective), as previously described (31). Briefly, stage 7-8 electroporated embryos were cultured under the microscope at 38 degrees in humid atmosphere. Two channels (GFP and brightfield), 3 fields of views, 10 Z levels were imaged every 6 minutes for each embryo (6 embryos per experiment). Images were stitched and pixels in focus were selected using Stack Focuser (ImageJ). X, Y drift was corrected using MultiStackReg adapted from TurboReg (ImageJ) (52). Image segmentation was done after background correction using Background Subtractor plugin (from the MOSAIC suite in ImageJ) and cell tracking was done using Particle Tracker 2D/3D plugin (ImageJ) (53). A reference point was defined for each frame at the last formed somite using manual tracking. Regions of interest were defined manually and their posterior movement was defined by manual tracking of the tailbud movement. Subtraction of the tissue movement was done by defining the average motions of cells in the region. Violin plots were generated on Prism 8 (Graphpad). MSD and distribution of angles were calculated and plotted with a Matlab routine. Angle distribution was calculated from trajectories, weighted with velocities and plotted as rosewind plot using Matlab.

### Statistical testing

Kolmorogov-Sminorv test was used to test for differences in angle distributions in Figure 5D and 5F. For all the other comparisons, unpaired Student test was used, P<0.05 *, P<0.001 **, P<0.0001 ***, P<0.00001 ****, P>0.05 non-significant (ns).

### Mathematical modeling

A number of 1100 progenitor cells, 1200 neural cells and 3200 PSM cells was initially distributed in their respective areas. Each cell type was endowed with its proliferation rate, i.e. 11.49 hours for progenitor cells, 10.83 hours for neural cells and 8.75 hours for PSM cells. Each cell I was characterized by a given ratio of Sox2/Bra, named R_I(t), with an assigned value from 0 to 2 (depicted as 0-2 in Figure 4A to match with biological ratios), and by a 2D position (x_I(t),y_I(t)), each of these variables being time-dependent. In the random model, an initial Sox2/Bra ratio value from 0.15 to 0.85 was randomly attributed to progenitor cells. At each time step, each cell updates its Sox2/Bra value through a first order ODE, using the function represented in Figure 5A (+ noise), and then updates its position (x,y), depending on the value of the ratio, by a biased/adapted random motion. Interaction properties between cells such as adhesion, maximum density, packing were implemented in the bias of the random motion, as detailed in the Supplementary Materials and Methods. Simulations focused on the posterior body (1 unit=150 micrometers). Cells movements in the most anterior region were blocked, considering this region, composed of somites and neuroepithelial cells, as a very dense area, and, similarly, cell passage to either side of the PSM was blocked, considering the lateral plate to be a solid structure.

## Acknowledgements

We thank Karine Guevorkian, Eric Théveneau, and Myriam Roussigné for critical reading of the manuscript. We thank Brice Ronsin and Stephanie Bosch and the CBI imaging facility and Marion Aguirrebengoa for help with statistics. We also thank members of the Pituello, Soula, Theveneau and Davy teams for suggestions and stimulating discussions during the project. This work has been funded by ANR (JC) and the ARC foundation grants.

---

**Supplemental Figure 1:**
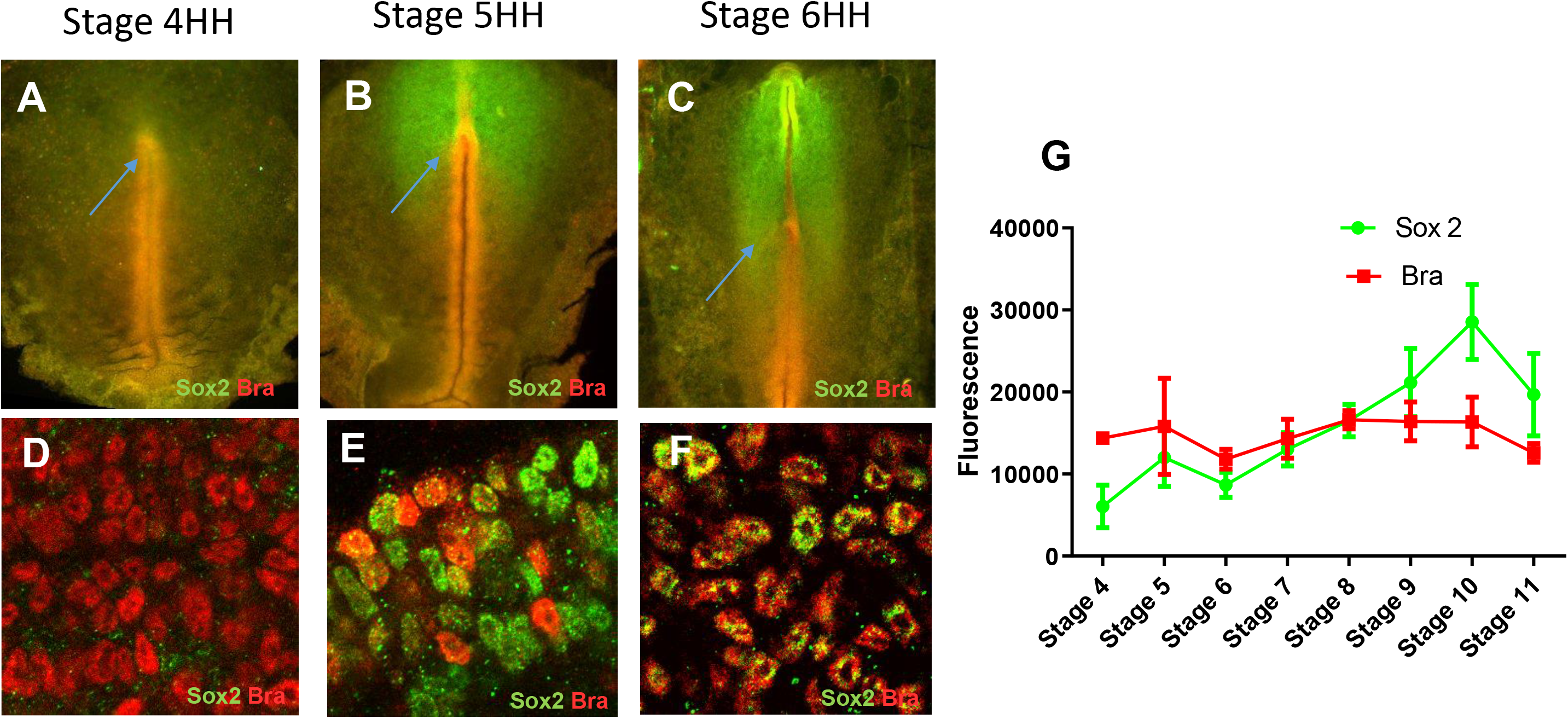
Sox2 and Bra protein co-expression and cell-to-cell heterogeneity start around stage HH5 in quail embryos. **A-F:** Double immunodetection of Sox2 and Bra viewed on whole-mount embryos (**A-C**) and at cell levels in the PZ (**D-F**) at stages HH4 (**A,D**), HH5 (**B,E**) and HH6 (**C,F**). **G**: Sox2 and Bra average levels in posterior progenitors at different stages of development.

**Supplemental Figure 2:**
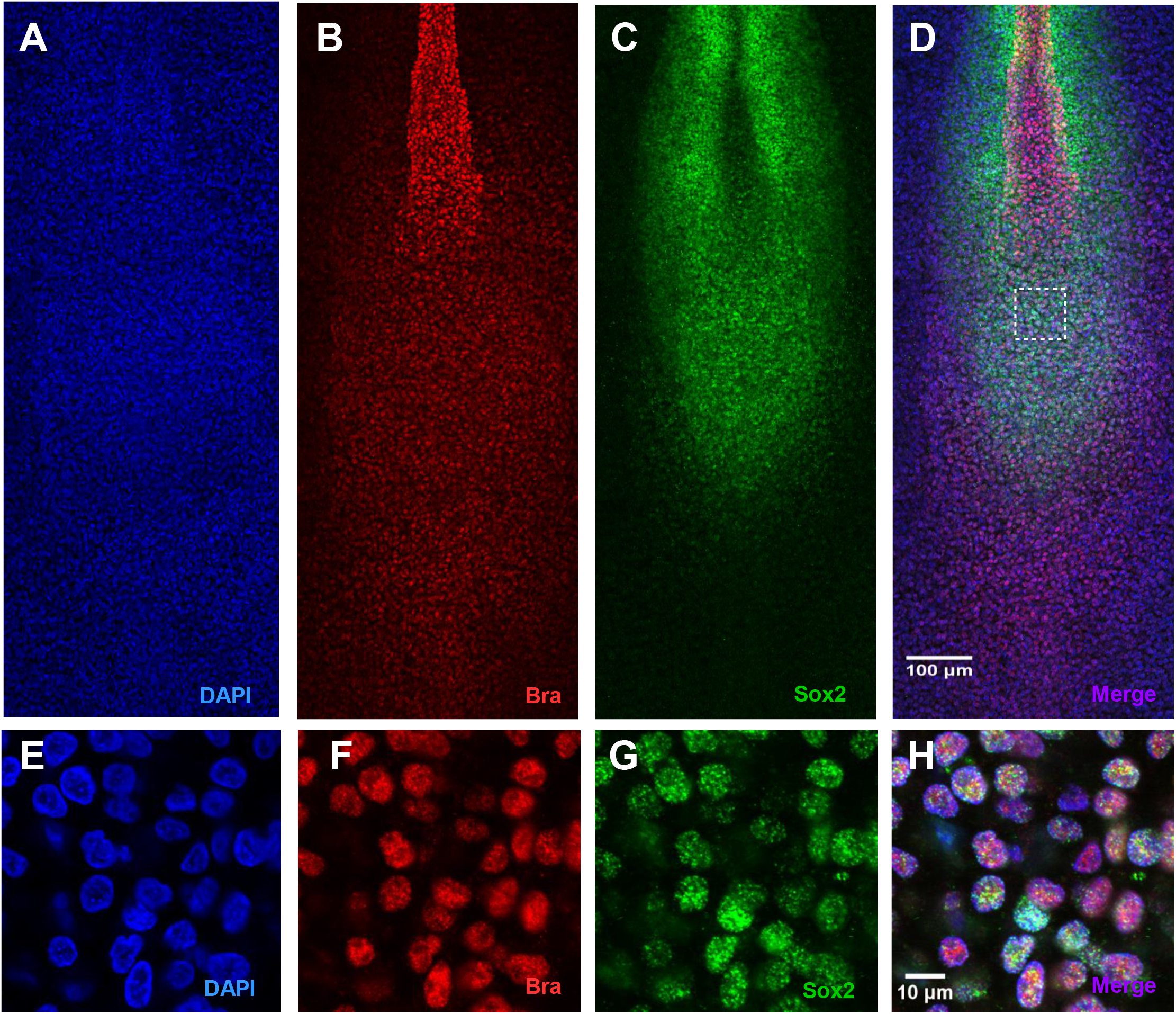
Chicken PZ cells co-express Sox2 and Bra proteins with a high degree of cell-to-cell heterogeneity. **A-H**: Double immunodetection of Bra and Sox2, counterstained with DAPI, on stage HH10 chicken embryo. Horizontal sets present successively the DAPI staining (**A, E**), the Bra (**B, F**) and Sox2 (**C, G**) signals and the merged images (**D, H**) on ventral views of the posterior region (**A-D**) and on a close up of the PZ (**E-H,** white square in **D**). Note variability in the Sox2 and Bra signal intensities in chicken PZ cells.

**Supplemental Figure 3:**
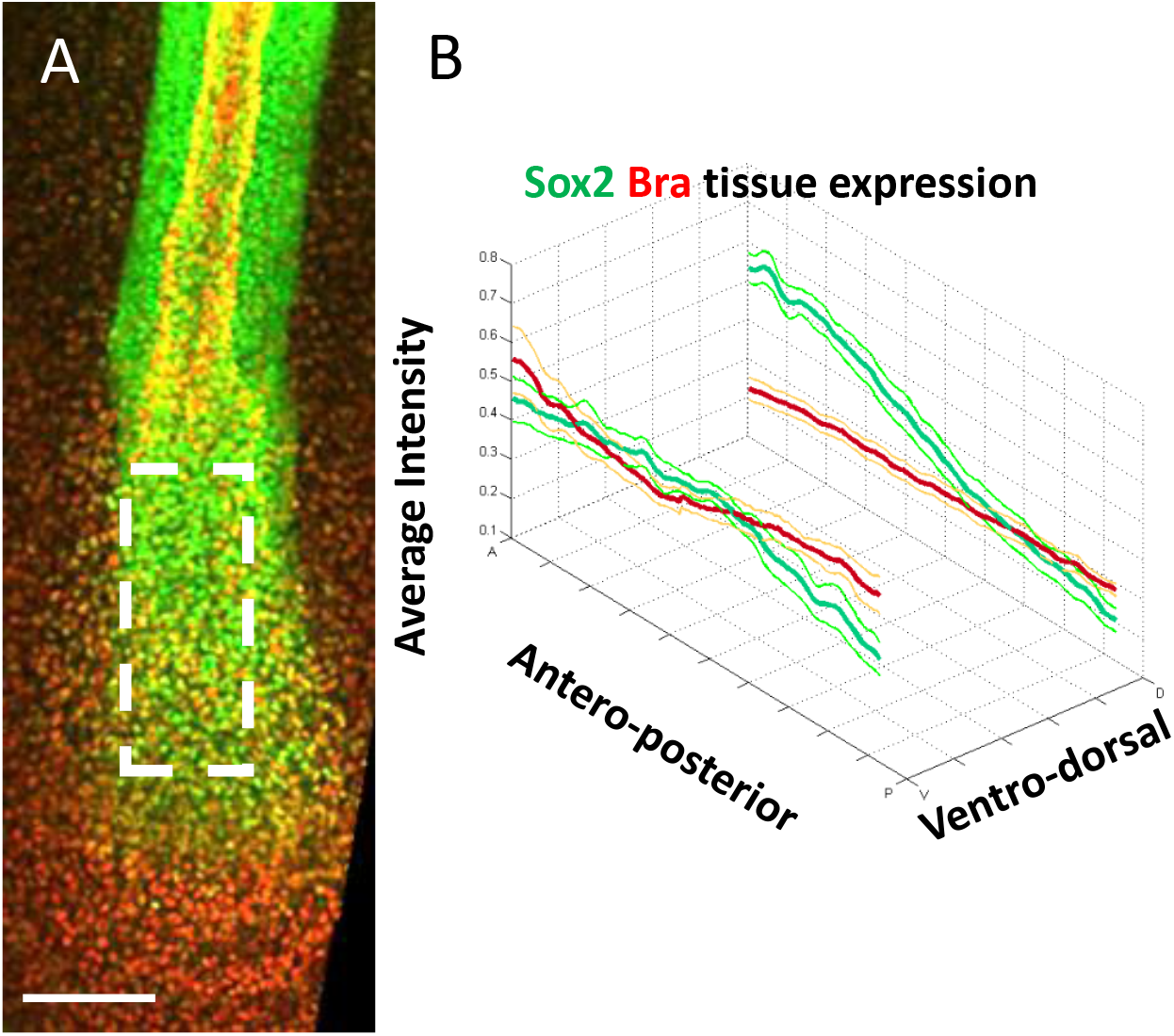
Sox2 and Bra expression patterns in PZ cells follow opposite gradients at the tissue level. **A**: Immunodetection of Sox2 (green) and Bra (red) viewed at the tissue scale in the caudal part of stage HH11 quail embryo. Dashed white rectangle delineates the PZ. **B**: Quantification of Sox2 (green) and Bra (red) proteins at different antero-posterior and dorso-ventral levels (n= 9 embryos, thick lines indicate average values, thin lines demarcate the SEM).

**Supplemental Figure 4:**
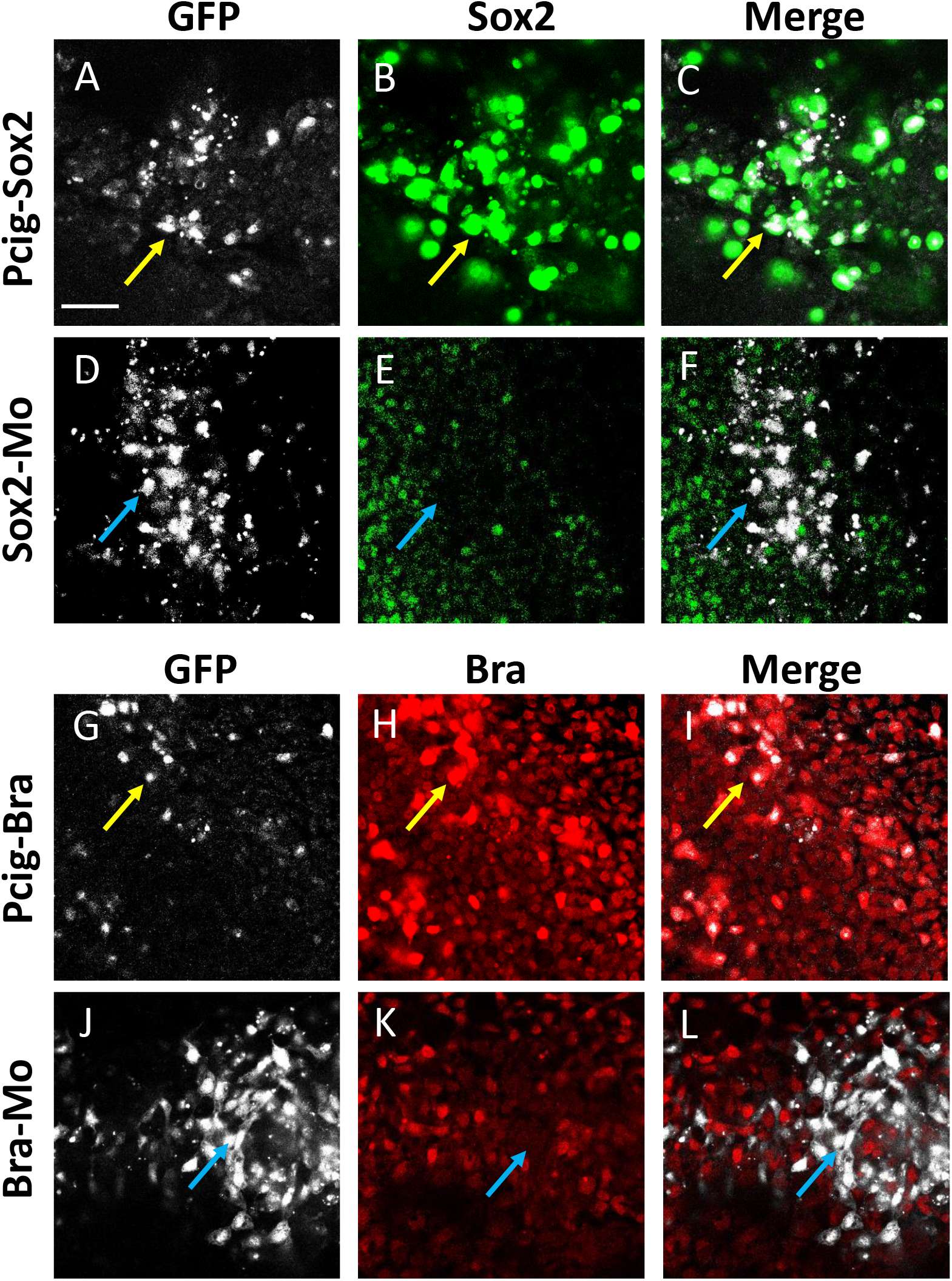
Efficient deregulation of Sox2 and Bra following electroporation of expression vectors and morpholinos. **A-F**: Immunodetection of Sox2 performed 7 hours after electroporation of the Pcig-Sox2 (**A-C**) and the Sox2-Mo (**D-E**). Horizontal sets show high magnification of the PZ and present successively the GFP signal (white, **A**, **D**), the Sox2 signal (green, **B**, **E**) and the merged images (**C, F**). Note co-localization of high Sox2 signal and GFP in Pcig-Sox2 transfected cells (yellow arrows in **A-C**) but not but not in Sox2-Mo transfected cells (blue arrows in **D-F**). **G-L**: Immunodetection of Bra performed 7 hours after electroporation of the Pcig-Bra (**G-I**) and the Bra-Mo (**J-L**). Horizontal sets show high magnification of the PZ and present successively the GFP signal (white, **G**, **J**), the Bra signal (red, **H**, **K**) and the merged images (**I**, **L**). Note co-localization of high Bra signal and GFP in Pcig-Bra transfected cells (yellow arrows in **G-I**) but not but not in Bra-Mo transfected cells (blue arrows in **J-L**) (scale bar is 50 μm).

**Supplemental Figure 5:**
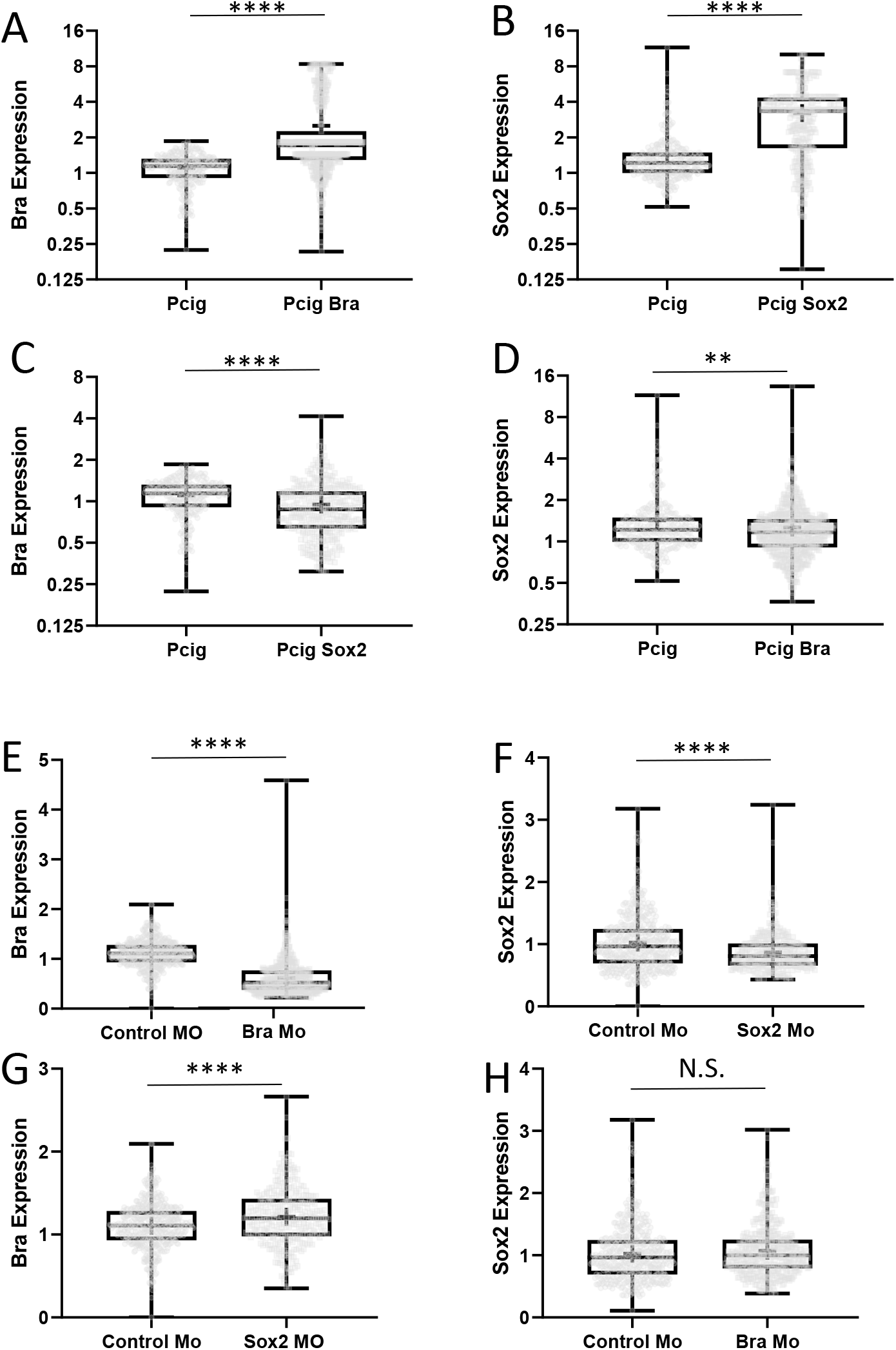
Quantification of Sox2 and Bra protein levels following electroporation of expression vectors and morpholinos. **A-D**: Quantification of Bra (**A**, **C**) and Sox2 (**B**, **D**) protein levels in cells electroporated with the Pcig-Bra (**A**, **D**) or the Pcig-Sox2 (**B**, **C**) expression vectors. Note the significant upregulation of Bra and Sox2 in cells electroporated with the Pcig-Bra and the Pcig-Sox2, respectively, compared to control cells (Pcig). Note also the significant down-regulation of Bra and Sox2 in cells electroporated with the Pcig-Sox2 and the Pcig-Bra, respectively. **E-H**: Quantification of Bra (**E**, **G**) and Sox2 (**F**, **H**) protein levels in cells electroporated with the Mo-Bra (**E**, **H**) or the Mo-Sox2 (**F**, **G**). Note the significant downregulation of Bra or Sox2 in cells electroporated with the Mo-Bra and the Mo-Sox2, respectively, compared to control cells (control Mo). Note also the significant upregulation of Bra in Mo-Sox2 electroporated cells but not that of Sox2 in Mo-Bra electroporated cells. Levels of protein expression are normalized to neighboring non-electroporated cells.

**Supplemental Figure 6:**
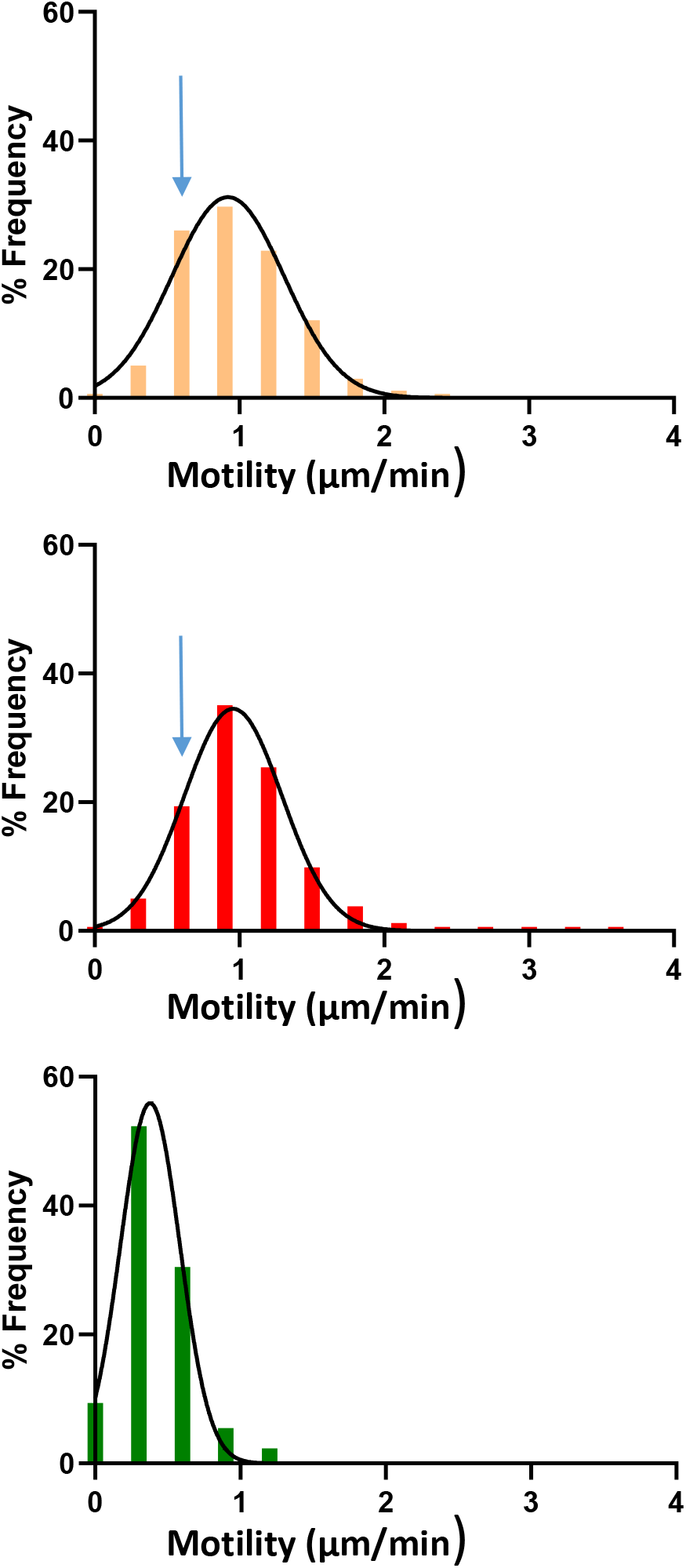
Distribution of motility frequencies within the PZ, the PSM and the neural tube. Representation of the distribution of frequency for different classes of speeds ranging from slow moving cells to fast moving cells corresponding to figure 3C corrected motility. Top panel (yellow) correspond to the PZ, middle panel (red) to the PSM and bottom panel (green) to the NT. Note that the distributions of speed are different between the three groups, the NT containing more slow cells than the two other tissues and the progenitor zone having slightly more slower cell than the PSM (higher histogram pointed by the blue arrow).

**Supplemental Figure 7:**
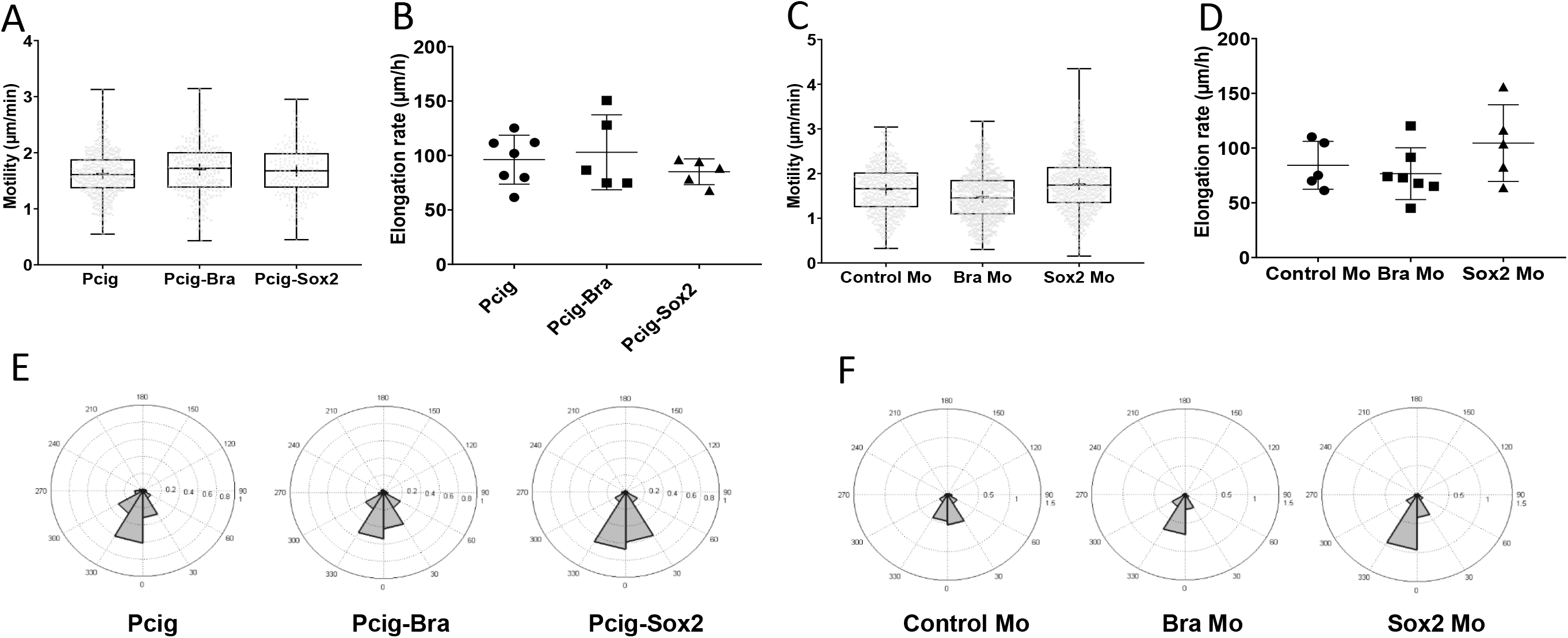
Effect of Sox2 and Bra on raw cell and tissue movements. Elongation (**A, C**), raw cell movements (**B, D**) and raw angle distribution (**E, F**) measurements for Sox2 and Bra over-expression (**A, B, E**) and down-regulation (**C, D, F**).

**Supplemental Figure 8:**
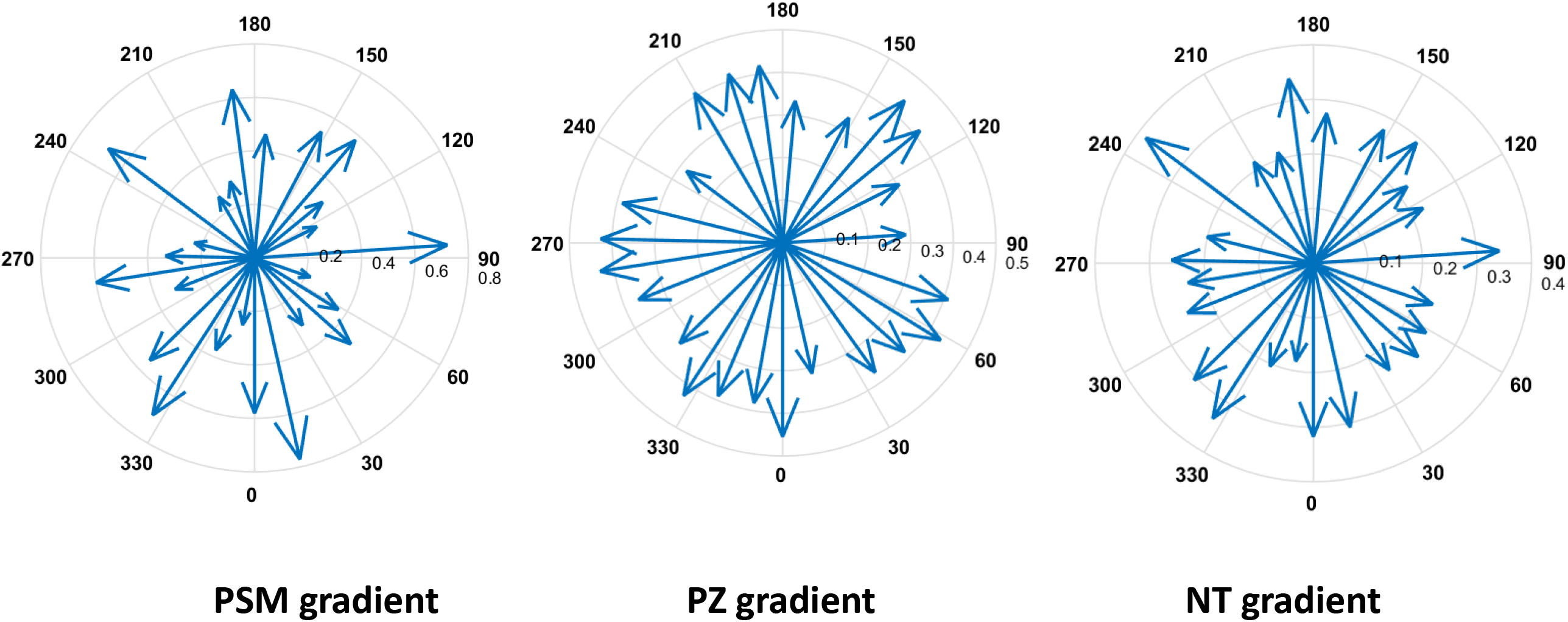
Analysis of directionality in the gradient model. Distribution of directionalities of migration within the PZ, the PSM and the NT of the gradient model.

**<u>Supplemental Movie 1</u>: Cellular migration in PZ, NT and PSM (corresponding to Figure 3).** H2B-GFP electroporated quail embryos. The movie shows a ventral view of the posterior region of a representative embryo (10x). GFP signal is in green, bright field in blue. Moving regions of interest are delineated as follow: yellow for the progenitor, red for the nascent paraxial mesoderm and green for the NT. Cell trajectories are marked and colored accordingly.

**<u>Supplemental Movie 2</u>: Posterior progenitor migration (corresponding to Figure 3).** H2B-GFP electroporated quail embryos. The movie shows a ventral view of the posterior region of a representative embryo (10x). The GFP signal is in white. The right movie show the raw signal and the left movie shows some example of cell tracking (yellow). Note that nuclei are often changing neighbors.

**<u>Supplemental Movie 3</u>: Mathematical simulation of posterior region behavior with Heterogeneous/random expression of Sox2 and Bra.**

**<u>Supplemental Movie 4</u>: Mathematical simulation with Heterogeneous expression of Sox2 and Bra: initial ratios set up for 50%.**

**<u>Supplemental Movie 5:</u> Mathematical simulation of Heterogeneous expression of Sox2 and Bra: Low Sox2/Bra (high Bra) case.**

**<u>Supplemental Movie 6</u>: Mathematical simulation of Heterogeneous expression of Sox2 and Bra: High Sox2/Bra (high Sox2).**

**<u>Supplemental Movie 7</u>: Progenitor migration in Sox2 and Bra overexpression experiments (Corresponding to Figure 5).** The movie shows ventral views of posterior regions of three representative embryos, focused on the PZ that has been set as a reference point (10x). GFP signal is in green, bright field is in blue. Cell trajectories are marked and colored in yellow. Bra overexpression is on the left, Sox2 overexpression on the right and control embryo is in the middle.

**<u>Supplemental Movie 8</u>: Progenitor migration in Sox2 and Bra Mo experiments (Corresponding to Figure 5).** The movie shows ventral views of posterior regions of three representative embryos, focused on the PZ that has been set as a reference point (10x). GFP signal is in green, bright field is in blue. Cell trajectories are marked and colored in yellow. Bra Mo is on the left, Sox2 Mo is on the right and control embryo is in the middle.

**<u>Supplemental Movie 9:</u> Mathematical simulation of posterior progenitor’s behavior: Graded expression of Sox2 and Bra case.**

**<u>Supplemental Movie 10:</u> Comparison of gradient (left) and heterogeneous (right) models.**

