## Supplementary Methods for "Cell-to-cell heterogeneity in Sox2 and Brachyury expression ratios guides progenitor destiny by controlling their motility."

### Supplemental Materials & Methods

#### 1 Part 1 : Image analysis

##### 1.1 Fixed tissue imaging

**Sox2 and Bra intensity profile from PZ to NT or PSM (Figure 1 G,G').** Cubes containing approximately 100 nuclei are defined with Imaris along a path going from the progenitor zone to the neural tube in one case, and from the progenitor zone to the PSM in the other case. The cubes are separated by 70 $\mu m$ distance approximately. The nuclei detection and data collection are done using the same methods as described previously for the immunodetection analysis. For each cube, the average of the nuclei normalised intensities is calculated and then reported to Graphpad to plot the intensity profile along the PZ-NT path or the PZ-PSM path.

**Immunodetection analysis and calculation of SOX2/Bra level of expression (Figure 1 H-J).** After the confocal acquisition, the volume acquired is analysed with Imaris. The acquisition is done with 3 colours channels: green channel is associated with the level of expression of Sox2, red channel is associated with Bra, and blue channel represents the DAPI staining. The regions of interest are drawn with the pointer directly on the confocal imaged volume represented on the Imaris viewer. The dimensions of the region of interest can be refined by entering the desired value in the dimension value fields. The "Spot Detection" function of Imaris is used on the blue channel (DAPI staining) to automatically detect fluorescent nuclei with a user-defined radius equal to the actual radius of a nucleus into the region of interest. The interesting data of the spot detected such as their identities, their position in the 3 dimensions of the space (X,Y,Z) and the level of intensity of the spot in the three channels (green, red, blue) are given by the "statistics" tab. The data are collected on a csv table, opened in an Excel results table where the data is organised by spot identity and their dimensions. The Excel table is read with a homemade Matlab routine where the intensities of the Bra (red channel) and Sox2 (green channel) are normalised with the intensity of DAPI (blue channel) to consider the loss of fluorescent signal due to the depth. This operation is done for each spot. Then, the routine calculates the ratio of the intensity of Sox2 on the intensity of Bra for each spot. The data are plotted with the "scatter" function of Matlab, in a 2D space representing the projection of the spots along 1 dimension depending on the orientation of the section used for the analysis. The spots are color-coded with their associated ratio value with a color bar going from red (low ratio, i.e high Bra intensity, low Sox2 intensity) to green (high ratio, i.e high Sox2 intensity, low Bra intensity). The scatter plots representing the Sox2/Bra ratio distribution are done with a Matlab code using the "scatter" function and an adapted version of functions "beeswarm" ([1]) for the distribution plot and "bplot" ([2]) for the whisker plot (10%-90% interval). The Sox2 and Bra expression distribution plots and the calculation of the coefficient of variation is done with Graphpad.

**Immunodetection analysis for Sox2 and Bra 3D intensity profiles (Supplemental Figure 3):** Raw confocal data are exported to ImageJ and automatically processed with a homemade ImageJ macro. First, a Gaussian filter is applied to the stack before using the binning tool giving a pixel with a value calculated from the average of 4 pixels (approximately $6\mu m$) in the x and y dimensions of the confocal stack. The macro process an average z-projection every 4 slices (approximately $8\mu m$) along the confocal stack with the plugin "Grouped Z Projector". Then, a region of interest is drawn along the antero-posterior axis with a 40-50 $\mu m$ width. For each pixel row of the region of interest (x dimension), the average of pixels value constituting the row is calculated, giving an intensity profile along the length (y dimension) of the region of interest. The operation is done for each average-projected slice. Then, an intensity profile is picked up every 4 average projected slices (approximately $32\mu m$), recorded in a CSV results table and read in Matlab to build a 3D plot of the intensity profiles with the plot3 function.

#### 1.2 Live imaging (Figure 3 and 5)

**Movie reconstruction.** The movie reconstruction has been done with an ImageJ/FIJI macro that automatizes several steps previously described ([3]). The movie reconstructed and processed is fully on-focus, aligned, and ready for the tracking phase.

**Tracking analysis.** The principle of the cell tracking method has been described ([4]). The cells are tracked with Particle 2D/3D plugin in FIJI using this principle, allowing the reconstruction of trajectories with spots detected frame by frame. The interesting data (trajectory identity, coordinates of trajectory spots along the time-lapse movie (X,Y,T)) are collected in a csv table results which is then read with a Matlab routine. Manual tracking of the last formed somite and the node are performed respectively to set the reference point and to get the displacement of the posterior area. The first steps of the automatized Matlab routine are to open the csv results table, to draw a reference line along the AP axis on the last frame of the movie using the "imline" function and to draw the region of interest (ROIs) on the first frame of the movie with "impoly" function. The ROIs will move along the movie accordingly to the manual tracking of the posterior area. Then, the trajectory spots contained in the ROIs are selected for analysis and their coordinates are corrected accordingly to the reference point (last formed somite). Spots are grouped by trajectory identity and calculations and plots are then performed.

**Motility and directionality calculation.** A displacement vector is computed for each cell, corresponding to the movement of a cell between two frames. A displacement vector forms a segment of a trajectory and its norm defines the distance travelled by a cell between these two frames (Fig.1). This vector is created using the spot's coordinates of a trajectory in the table results, for every time interval along the trajectory. From this vector, motility and directionality are calculated. The motility corresponds to the norm of the displacement vector $\Delta\vec{r}$ of a cell between two frames divided by the time interval between two frames $\Delta t$.

$$\Delta r(t+\Delta t) = ||\Delta\vec{r}(t+dt)|| = \sqrt{(x(t+\Delta t)-x(t))^2+(y(t+\Delta t)-y(t))^2} \quad (1)$$

where x and y are the coordinates of the displacement vector.

$$motility(t+\Delta t) = \frac{\Delta r(t+\Delta t)}{\Delta t} \quad (2)$$

The directionality is the property of being directional or maintaining a direction.

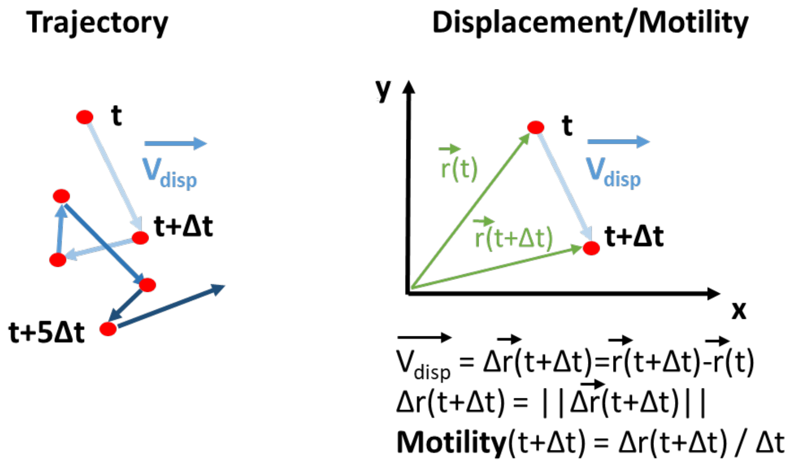


Figure 1: Trajectory and displacement/ motility definition and calculation for tracking analysis

In physics, this term is used to describe the preferential sensibility of a sensor or a broadcast system to a direction rather than others. Here, this term is used to describe the directional amplitude of a cell movement, taking into account its motility. First, the displacement vector ($\vec{v}$) between 2 frames is first compared to the reference line drawn previously ( $\overrightarrow{v_{ref}}$) as shown in Fig.2. The angle between the displacement vector and the reference line is calculated for each time interval along the trajectory with the following formula using the cross product and the dot product between the two vectors :

$$Angle(t) = \tan^{-1} \frac{||\vec{v}(t) \wedge \overrightarrow{v_{ref}}||}{\vec{v}(t) \cdot \overrightarrow{v_{ref}}} \tag{3}$$

The angle calculation is done for all the trajectories inside a ROI, and the displacement vectors are then allocated in twelve intervals ($int_\Theta$) from 0° to 360° (30° intervals) depending on their angle values. The number of vectors allocated to an interval ($n(\overrightarrow{v_{disp}}(int_\Theta))$) is divided by the total number of displacement vectors inside the ROI ($n_{total}(\overrightarrow{v_{disp}})$) to get the direction proportion of the displacement vectors $\tau(int_\Theta)$ in the interval.

$$\tau(int_\Theta) = \frac{n(\overrightarrow{v_{disp}}(int_\Theta))}{n_{total}(\overrightarrow{v_{disp}})} \tag{4}$$

This proportion is weighted by the average motility $< mot(\overrightarrow{v_{disp}}(int_\Theta)) >$ calculated from displacement vectors whose angle is contained in this angle interval, giving the directional amplitude $A(int_\Theta)$ of the cell movement in this direction. The calculation is done for every angle interval.

$$A(int_\Theta) = \tau(int_\Theta) \cdot < mot(\overrightarrow{v_{disp}}(int_\Theta)) > \tag{5}$$

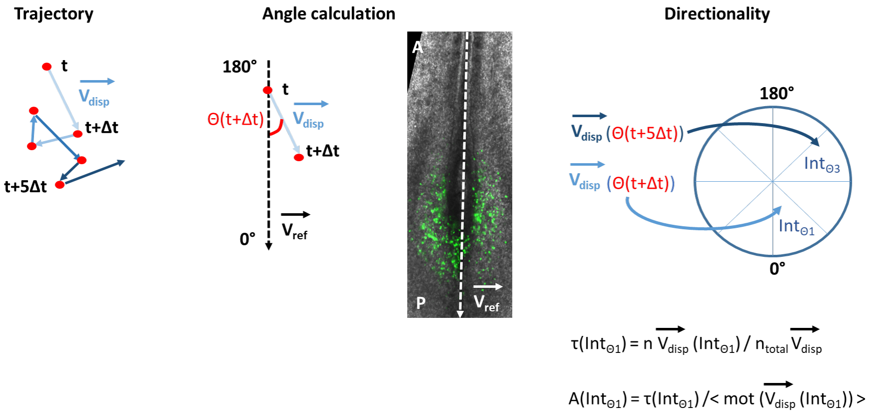


Figure 2: Angle and directionality calculation

**MSD calculation.** The mean square displacement (MSD) measures the deviation of the position of a particle with respect to a reference position over time. The MSD is used to analyse the mode of displacement of a particle, and is commonly applied in biophysics for studying cells movements in vivo. Here, the MSD is calculated for each cell trajectory using the spots coordinates from the table results within a defined time interval $\tau$.

$$MSD(\tau) =< \Delta r^2(\tau) >=< [r(t+\tau) - r(t)]^2 > \tag{6}$$

where$r(t)$ is the position of the particle at time $t$, and $\tau$ is the lag time between the two positions taken by the particle used to calculate the displacement $\Delta r(\tau) = r(t+\tau) - r(t)$. The MSD is then averaged for all the particles contained in a ROI.

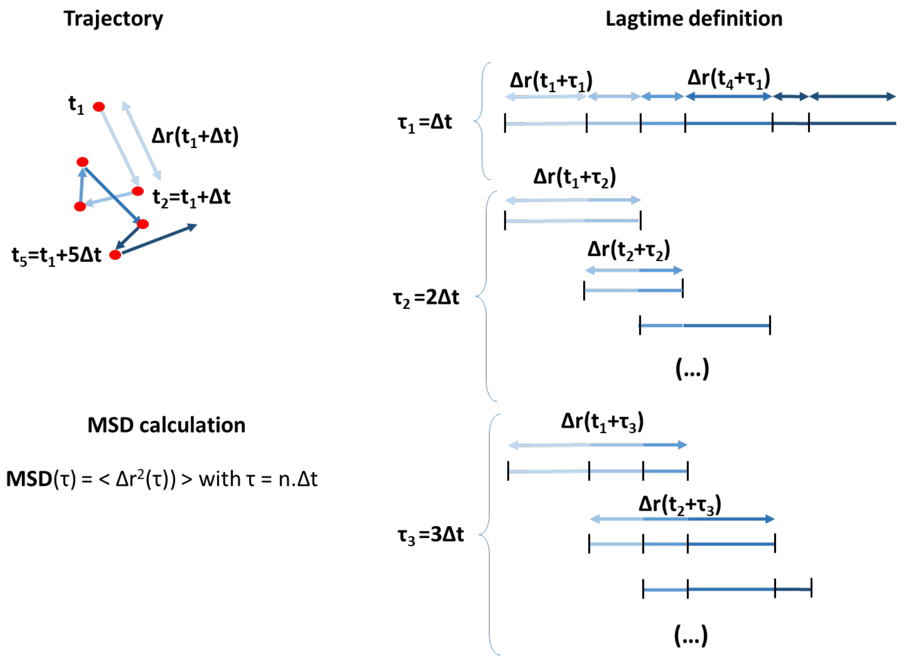


Figure 3: Lagtime definition and MSD calculation

**Average vector subtraction.** According to (20) the average movement of electroporated cells in the ROIs corresponds to the extracellular matrix movements. For each ROIs, the average velocity vector is defined for each frame by calculating the average velocity and the average direction (i.e the average angle of the displacement vectors with the reference line) with the coordinates of all the trajectories spots inside the ROI between this frame and the previous one. The average velocity vector is subtracted frame by frame to give the corrected coordinates of all the trajectories inside the ROI, and the corrected motility, MSD and directionality are calculated from these corrected data using the same formulas as previously.

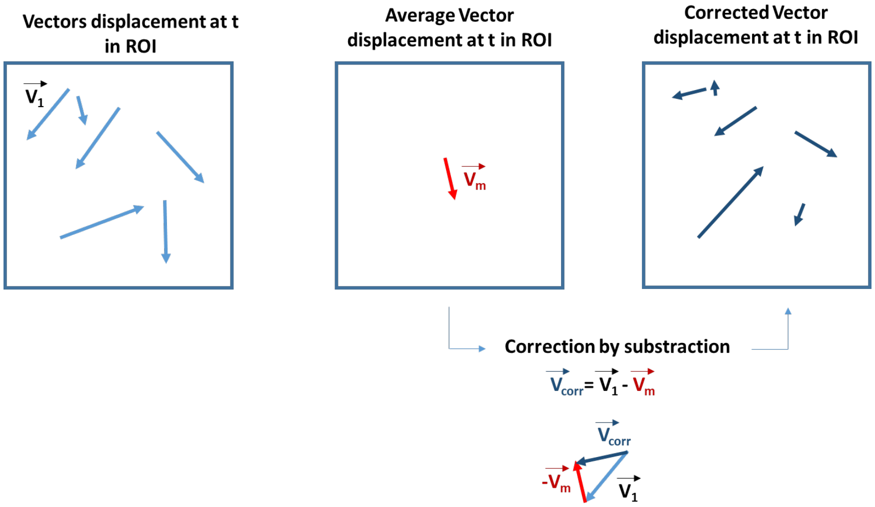


Figure 4: Average vector substraction for vector corrrection

**Save data.** The data are collected and are saved in tables in the ".mat" format in a specific folder corresponding to each embryo.

**Plots.** The ".mat" files containing the data for plots are read with a Matlab routine which collects the data per variables (motility, MSD, directionality), conditions and ROIs for all the embryos. The MSD is averaged for all embryos per lag time and the directionalities are averaged for all embryos per angle intervals determined previously. For the MSD and the motility distributions, table results are opened with Matlab and then transfered to Graphpad. The rose wind plots are made in Matlab according to the number of angle intervals and using the "rose and polar" functions from Matlab, showing the distribution of cells directionality in angle bins (here, 12 angle bins).

### 2 Part 2 : Mathematical Modeling

In this section we will exhibit the details of the modeling choices. We develop two agent based models in 2D to better understand how the regulation of cell motility by specification factors Sox2 and Bra can influence progenitor behaviors during axis elongation. The two models are: the heterogeneous model (Sox2/Bra levels are randomly distributed among progenitors), and the gradient model (Sox2/Bra levels are patterned in gradient among progenitors).

#### 2.1 The heterogeneous model

**Main variables of the model:** In this model the main variables are the cell type (defined by its Sox2/Bra level, details below), which defines its color in the simulation, and the cell position (random motion). Each of these variables is time-dependent.

##### 2.1.1 Differenciation and evolution of Sox2/Bra levels

Each cell $i$ is characterized by its ratio $R_i(t)$ defining the $\frac{Sox2}{Bra+Sox2}$ dynamics: A ratio $R_i(t) = 0$ corresponds to a PSM cell (a red cell in the simulation), and a cell of ratio $R_i(t) = 1$ corresponds to a neural cell (a green cell in the simulation). Finally, a cell with a ratio $0 < R_i(t) < 1$ is a progenitor cell (different shades of yellow in the simulation), that has not yet differentiated. To model heterogeneity and oscillations of transcription factors $R_i(t)$ of a cell $i$:

$$d_t R_i(t) = \Bigg(100(R_i(t)-0.2)(R_i(t)-0.5)(R_i(t)-0.8) + k_r dB_1(t)\Bigg)\mathbb{1}_{R_i\in(0,1)} \quad (7)$$

with $dB_1(t)$ the derivative of the Brownian motion associated to the ratio, it represents the various signals a cell receives, and affecting the expression levels of Sox2 and Bra, thus leading it to either cell fate: NT or PSM, corresponding to a ratio of 1 or 0 respectively. Moreover, $k_r$ is a constant representing the intensity of the signal, it is related to the specification rate of the cells. Its value is such that the number of progenitors remains constant throughout the phases of development we are modeling, then in a way to keep the balance between the specification of the cells into either fates and their proliferation (see below for details on the proliferation). Finally, a cell with a ratio $0 < R_i(t) < 0.2$ is called a pre-mesodermal cell, whereas a cell with a ratio $0.8 < R_i(t) < 1$ is called a pre-neural cell.

Remark 1: In practice for the numerical simulations we work discretely in time thus we replace the Brownian motion with a uniform variable $U(t) \in [-1; 1]$ at each time step.

Remark 2: Ignoring the noise, the deterministic function governing the evolution of the ratio has 5 equilinriums at $0, 0.2, 0.5, 0.8$, and 1.

Remark 3: The dynamics we chose for the ratio : $\frac{Sox2}{Bra+Sox2}$ is a simple example. In fact one can imagine a more general quantity in $[0; 1]$ describing the dynamics.

###### 2.1.2 Diffusion process

From litterature ([5]) and the present analysis (Figure 3) we know that cells in the progenitor zone, in the PSM and in the neural tube have a non-directional motility. Thus, one way to model cell motility is using a random unbiased walk (diffusion process) whose intensity depends on the cell's ratio $R_i$. The ODEs read:

$$d_t x_i(t) = k_x dB_2(t) \times \mathcal{V}(R_i(t)), \tag{8}$$

$$d_t y_i(t) = k_y dB_3(t) \times \mathcal{V}(R_i(t)), \tag{9}$$

with $dB_2(t)$ and $dB_3(t)$ the derivatives of the Brownian motions associated to the position (in $x$ and in $y$) of the cell. They represent the noise of the random walk affecting the cell's movements.
Remark : In practice, for the numerical simulations we work in a discrete environment in time, thus we replace the Brownian motions with uniform variables $v_x(t), v_y(t) \in [-1; 1]$ respectively replacing $dB_2(t)$ and $dB_3(t)$.
Furthermore, the variables $k_x$ and $k_y$ represent the intensity of the diffusion. The values of these two variables were chosen to fit biological data (figure 3 C), in accordance with our scale (here we took 1u =150 $\mu m$). Furthermore, the velocity function $\mathcal{V}(R_i(t))$ has the following form:

$$\mathcal{V}(R_i(t)) = \begin{cases} 1 + \beta(\tilde{R} - R_i(t))^2 & \text{if } 0 \leq R_i(t) \leq \tilde{R} \\ 1 & \text{if } \tilde{R} \leq R_i(t) \leq R^* \\ 1 - \alpha(R^\star - R_i(t))^2 & \text{if } R^* \leq R_i(t) \leq 1 \end{cases}$$

with $\alpha$ and $\beta$ two positive constants: $\alpha = \frac{1-0.65}{(R^\star-1)^2}$ and $\beta = \frac{1-0.8}{\tilde{R}^2}$.
The velocity of each cell $i$ depends on its Sox2/Bra ratio: it is equal to 1.2 ($\mathcal{V}(R_i(t))$=1.2) for PSM cells, and equal to 0.65 ($\mathcal{V}(R_i(t))$=0.65) for neural cells. As for cells in the PZ with a ratio above the neural threshold $0 < R^\star < 1$ (pre-neural cells), they will have a velocity between 0.65 and 1, and cells below the mesodermal threshold $0 < \tilde{R} < 1$ (pre-mesordermal cells) , they will have a decreasing velocity from 1.2 to 1. Cells with a ratio in between the pre-neural and the pre-mesordermal thresholds will have a constant velocity equal to 1.

###### 2.1.3 Proliferation

Each cell type is endowed with its proliferation rate: 11.49 hours for the progenitor cells, 10.83 hours for the neural cells and 8.75 hours for PSM cells ([6]). Then each cell, depending on its type, has a probability to proliferate given by $b_{PSM}$ for PSM cells, $b_{NT}$ for NT cells and $b_{PZ}$ for progenitors:

$$b_{PSM} = \frac{1}{8.75}, \quad b_{NT} = \frac{1}{10.83}, \quad b_{PZ} = \frac{1}{11.49}. \tag{10}$$

When each mother cell proliferates, it gives rise to one daughter cell, which inherites instantly its position and its ratio. The daughter cell is then integrated in the model, and varies its position through the ODEs (8) and (9), and its ratio through (7). Depending on its type (PSM, NT, PZ) it is added to the total cell number of its corresponding population.

###### 2.1.4 Biophysical properties and cell-cell interactions:

The system is confined between two horizontal lines, at $x = 1.5$ and $x = -1.5$, representing the lateral plate which, based on its higher cellular density ([5],[6]) can be assumed to act as a physical barrier on either side of the PSM. The system is also limited from the most anterior part, at $y = 2$, as we consider everything above that line to be of high density/epithelial (somites and anterior neural tube). Recall the scale we chose: $1u = 150\mu m$ ($u$= graph unit), and here we represent a portion of the posterior body (the most posterior), considering that a portion of the PSM and the NT have already been formed (further details in the pargraph Initial condition).
Cells are endowed with properties allowing them to interact with each other. Indeed cells with high Sox2 adhere, more or less depending on their level of Sox2. The adhesion dynamic we chose is to ask cells to redirect their jump (in $x$ and in $y$) as long as their immediate neighborhood $\epsilon$ (a ball of radius $\epsilon$ representing $15\mu m$) does not contain a certain number of neighboring cells of the same type. As previously mentioned that number depends on the level of Sox2 in each cell, then cells of the neural tube adhere the most. This is a natural assumption as we know that the neural tube is an epithelium, where cells are closely packed. Moreover, cells tend to move away from foreign high densities within a certain neighborhood (a ball of radius $2\epsilon$ representing $30\mu m$). In particular, cells with various levels of Bra flee densities of all foreign cells. Furthermore, to keep well defined physical boundaries between the tissues we also ask neural cells to move away from high densities of PSM cells. Using these interaction rules we created a model that presents self-organizing phenomena, with physical boundaries between the tissues.

**Initial condition:** We consider the embryo to be at a stage where a portion of the NT and the PSM have already been formed (anteriorly), for example at stage HH8. We distribute 1200 neural cells inside the square $[-0.5; 0.5] \times [0; 2]$, with an initial ratio of 1. As these cells are already differentiated into their neural fate, their ratio is not updated at each time step. We distribute 3200 PSM cells, which corresponds to 1600 from either side of the already formed neural tube, inside $[-1.5; -0.5] \cup [0.5; 1.5] \times [-2.5; 2]$, with an initial ratio of 0. These cells also don't update their ratio as they have already engaged in their mesodermal fate. Finally, we distribute 1100 progenitor cells inside the square $[-0.5; 0.5] \times [-1.5; 0]$. Each progenitor cell $i$ is attributed a random initial ratio in $[0.15; 0.85]$.
Remark: By definition of the ratio and by equation (7), we can see that progenitor cells with a ratio less than 0.2 are predestined to a mesodermal fate, and cells with a ratio higher than 0.8 are predestined to a neural fate. These predestined cells account for 14% of the total number of progenitor cells initially. The presence of these specified cells is justified by the fact that we have noticed

the presence of progenitors having Sox/Bra levels as high or as low as NT or PSM cells in the biological system ( Figure 1 I)

**List of variables and their values:** $k_r = 4.68$
$\epsilon = 0.1$, corresponding to $15\mu m$.
maxTN=5, representing the maximal density (number of cells in a neighborhood of size $2\epsilon$) a neural cell can withhold before fleeing the density.
maxPSM=4, representing the maximal density (number of cells in a neighborhood of size $2\epsilon$) a PSM cell can withhold before fleeing the density.
maxPZ=4, representing the maximal density (number of cells in a neighborhood of size $2\epsilon$) a progenitor cell can withhold before fleeing the density.
$\Delta t = 0.01$, the time step of the scheme, representing 6 seconds (or 0.00167 hours) of real biological development.
Tmax=60, representing 10 hours of real biological development.
$k_x = 7.5$
$k_y = 7.5$
$R^\star = 0.8$
$\tilde{R} = 0.2$
Remark: One should note that the frame rate of the live imaging is 1 frame per 6 minutes. Thus we use the same rate for the simulations: 1 frame per 6 minutes, corresponding to 1 frame per $60\Delta t$.

###### 2.1.5 Case 1: high Sox2

To simulate the high Sox2 case, the only parameter we change is the noise $U(t)$ in the ODE of the ratio $R_i(t)$. In fact, instead of having a uniform variable in $[-1; 1]$, allowing cells to have an equal probability of differentiating to either PSM or neural cells, we shift the noise and get a uniform variable in $[-0.97; 1.03]$. By doing that, cells have a higher probability to differentiate towards a neural ratio ($R_i(t) = 1$).

###### 2.1.6 Case 2: high Bra

To simulate the high Sox2 case, the only parameter we change is the noise $U(t)$ in the ODE of the ratio $R_i(t)$. In fact, instead of having a uniform variable in $[-1; 1]$, allowing cells to have an equal probability of differentiating to either PSM or neural cells, we shift the noise and get a uniform variable in $[-1.03; 0.97]$. By doing that, cells have a higher probability to differentiate towards a PSM ratio ($R_i(t) = 0$).

##### 2.2 The gradient model

The purpose of this model is to impose a gradient-like structure to the progenitor zone. More specifically, our aim is to have cells expressing high Sox2 in the most anterior progenitor zone, and cells expressing high Bra in the most posterior progenitor zone, while keeping the pool of progenitors. To do so, we divide our progenitor zone into 8 subdivisions in the direction of the y-axis, such that each subdivision contains the same number of cells. Cells in the subdivisions 1 to 3 will be subjected to a signal coding for an over-expression of Sox2 (biased noise similar to the case 1), and cells in the subdivisions 6 to 8 will be subjected to a signal coding for an over-expression of Bra (biased noise similar to the case 2), while cells in the subdivisions 4 and 5 will be subjected to a weak signal (noise) to keep the ratio of these cells near the equilibrium state 0.5, which will allow them to proliferate and keep the progenitor pool. Then, calling $f(R_i(t)) := 100(R_i(t)-0.2)(R_i(t)-0.5)(R_i(t)-0.8)$ the ODE on the ratio reads:

$$d_t R_i(t) = \begin{cases} f(R_i(t)) + k_r dB_a(t) & \text{if } subdiv(1) \leq y_i(t) \leq subdiv(3) \\ f(R_i(t)) + k_r dB_b(t) & \text{if } subdiv(4) \leq y_i(t) \leq subdiv(5) \\ f(R_i(t)) + k_r dB_c(t) & \text{if } subdiv(6) \leq y_i(t) \leq subdiv(8) \end{cases}$$

with $dB_a(t), dB_b(t), dB_c(t)$ the derivatives of the Brownian motion of the cell in each division of the zone, respectively replaced by $U_1(t), U_2(t), U_3(t)$ in the numerical simulations, with $U_1(t)$ a uniform random variable in $[-0.9; 1.1]$ corresponding to a high Sox2 signal in this region, $U_3(t)$ a uniform random variable in $[-1.1; 0.9]$ corresponding to a high Bra signal in this region, and $U_2(t)$ a uniform random variable in $[-0.1; 0.1]$ corresponding to a weak signal in this region where ratios will vary slightly, thus inhibiting specification and favoring the maintenance of the progenitor pool. Thus, at each time step, cells will evaluate their position, and update their ratio accordingly. We also note that at each time step the gradient is updated to the new progenitor zone (that has elongated).
Remark: We do not change the velocity function $\mathcal{V}(R_i)$, nor the interaction between cells nor the biophysical properties of the model. However, the initial distribution of the progenitor cells is changed to be gradient patterned (details in the paragraph Initial condition).

**Initial condition:** The disposition of the tissues PSM, TN, and PZ remains the same as the heterogeneous model. The initial distribution of the PSM and NT cells is also unchanged. However, we change the initial distribution of the PZ cells from a random one, to a gradient distribution as the following:
we distribute 1100 progenitor cells inside the square $[-0.5; 0.5] \times [-1.5; 0]$. Each progenitor cell $i$ is attributed an initial ratio in $[0.15; 0.85]$, depending on its position in the progenitor zone: cells with the highest ratio (closer to 1) will be placed anteriorly, whereas cells with the lowest ratio (closer to 0) will be placed posteriorly. Using a linear function in $y$ the ratios are initially distributed as the following:

$$R_i(t=0) = -\frac{0.15 - 0.85}{1.5} y_i(t=0) + 0.85 \tag{11}$$

Remark: By definition of the ratio, and equation (7), and what we just described, we can see that progenitor cells with a ratio less than 0.2, predestined to a mesodermal fate, are now in the most posterior region of the PZ, and cells with a ratio higher than 0.8, predestined to a neural fate, are now placed in the most anterior part of the PZ.

**List of variables and their values:** We tried to keep most variables equal to the ones in the heterogeneous model, however some of them are not comparable then we could change their values to meet the model hypotheses.
$k_r = 4.03$
$\epsilon = 0.1$, corresponding to $15\mu m$.
maxTN=5, representing the maximal density (number of cells in a neighborhood of size $\epsilon$) a neural cell can withhold before fleeing the density.
maxPSM=4, representing the maximal density (number of cells in a neighborhood of size $\epsilon$) a PSM cell can withhold before fleeing the density.
maxPZ=4, representing the maximal density (number of cells in a neighborhood of size $\epsilon$) a progenitor cell can withhold before fleeing the density.
$\Delta t = 0.01$, the time step of the scheme, representing 6 seconds (or 0.00167 hours) of real biological development.
Tmax=60, representing 10 hours of real biological development.
$k_x = 7.5$
$k_y = 7.5$
$R^\star = 0.8$
$\tilde{R} = 0.2$
Remark: One should note that the frame rate of the live imaging is 1 frame per 6 minutes. Thus we use the same rate for the simulations: 1 frame per 6 minutes, corresponding to 1 frame per $60\Delta t$.

### 3 Figures

In the following paragraph, we present the details of the computations of the violin plots and the angle plots. We used this approach to validate the model and compare its results to those of the live imaging analysis.
We do the same computations for the heterogeneous model and for the gradient model.

**Cell tracking for both models:** To generate the violin plots (figure 4 C,G, figure 6 C), we save the cells' positions every 6 minutes (in real biological time, to correspond to the time points of the live imaging), this corresponds to $60\Delta t$ in simulation time with our choice of $\Delta t$. To compute the cell velocity in each tissue, we consider a region of size $1u \times 1.5u$ in the PZ and the NT, and $0.8u \times 2u$ in the PSM. Then, in each tissue, we compute the velocity of each trajectory for every cell, between the time point $t$ and $t + 6$minutes, starting from $t = 0$ to $t = 10$ hours. We only consider the cells up to the time when they exit the regions.
To plot the distribution of the angles in each tissue (figure 4 D, supp figure 8), for every cell type we consider each cell (in the regions previously drawn in

each tissue), and compute the angle between the trajectory and the reference vector pointing downwards (in the direction of the elongation) of each trajectory, between the time point $t$ and $t + 6$minutes . This angle will fall into one of 24 bins (dividing the 360° disk into 24 subdivisions). Finally we compute the mean velocity in every bin. We then multiply each mean velocity, in each bin, by the proportion of trajectories in that bin, this gives the length of the arrow plotted in each bin.

**Tracking the shape of the progenitor zone for both models:** We upload the simulation generated by Matlab to the software ImageJ and manually track the most anterior and most posterior point of the progenitor zone (per frame). This gives a track of the length of the progenitor zone through time. For the width, as the progenitor zone is moving and changing shapes, we choose to track the width of the mid-progenitor zone throughout the simulation (tracking the mid-left and mid-right points). We then plot the rectangles hereby generated (length × width) for initial time and final time.
We use the manual tracking of the most posterior point of the progenitor zone to compute the elongation rate.

**Mean square displacement for both models:** To compute the mean square displacement of progenitor cells we use the following formula:

$$MSD(t) = \frac{1}{N}\sum_{i=1}^{N}\left((x_i(t) - x_i(0))^2 + (y_i(t) - y_i(0))^2\right) \qquad (12)$$

with $N$ the number of cells considered. Here we consider all the progenitor cells up to the time when they differentiate.

**Numerical scheme for both models:** For the heterogeneous model and the gradient model, we use a finite-difference scheme to discretize the ODEs. The code was done using Matlab R2020a.
